# Highly Efficient Hypothesis Testing Methods for Regression-type Tests with Correlated Observations and Heterogeneous Variance Structure

**DOI:** 10.1101/552331

**Authors:** Yun Zhang, Gautam Bandyopadhyay, David J. Topham, Ann R. Falsey, Xing Qiu

**Affiliations:** J Craig Venter Institute, 4120 Capricorn Lane, La Jolla, CA 92037, United States; Department of Surgery, University of Rochester, 601 Elmwood Ave, Rochester, NY 14642, United States; Department of Microbiology and Immunology, University of Rochester, 601 Elmwood Ave, Rochester, NY 14642, United States; Department of Medicine, University of Rochester, 601 Elmwood Ave, Rochester, NY 14642, United States

**Author notes:** Correspondence, Department of Biostatistics and Computational Biology, University of Rochester, 601 Elmwood Ave, Rochester, NY 14642, United States.

**Keywords:** Hypothesis testing, Matrix decomposition, Orthogonal transformation, RNA-seq; Rotated test

## Abstract

**Background:** For many practical hypothesis testing (H-T) applications, the data are correlated and/or with heterogeneous variance structure. The regression *t*-test for weighted linear mixed-effects regression (LMER) is a legitimate choice because it accounts for complex covariance structure; however, high computational costs and occasional convergence issues make it impractical for analyzing high-throughput data. In this paper, we propose computationally efficient parametric and semiparametric tests based on a set of specialized matrix techniques dubbed as the PB-transformation. The PB-transformation has two advantages: 1. The PB-transformed data will have a scalar variance-covariance matrix. 2. The original H-T problem will be reduced to an equivalent one-sample H-T problem. The transformed problem can then be approached by either the one-sample Students *t*-test or Wilcoxon signed rank test.

**Results:** In simulation studies, the proposed methods outperform commonly used alternative methods under both normal and double exponential distributions. In particular, the PB-transformed *t*-test produces notably better results than the weighted LMER test, especially in the high correlation case, using only a small fraction of computational cost (3 versus 933 seconds). We apply these two methods to a set of RNA-seq gene expression data collected in a breast cancer study. Pathway analyses show that the PB-transformed *t*-test reveals more biologically relevant findings in relation to breast cancer than the weighted LMER test․.

**Conclusions:** As fast and numerically stable replacements for the weighted LMER test, the PB-transformed tests are especially suitable for “messy” high-throughput data that include both independent and matched/repeated samples. By using our method, the practitioners no longer have to choose between using partial data (applying paired tests to only the matched samples) or ignoring the correlation in the data (applying two sample tests to data with some correlated samples).

## 1 Background

Modern statistical applications are typically characterized by three major challenges: (a) high-dimensionality; (b) heterogeneous variability of the data; and (c) correlation among observations. For example, numerous data sets are routinely produced by high-throughput technologies, such as microarray and next-generation sequencing, and it has become a common practice to investigate tens of thousands of hypotheses simultaneously for those data. When the classical *i.i.d.* assumption is met, the computational issue associated with high-dimensional hypothesis testing (hereinafter, H-T) problem is relatively easy to solve. As proof, R package `genefilter` (Gentleman et al., 2017) implements a vectorized computation of the Student’s *t*-test which is more than a hundred times faster than the stock R function `t.test()`. However, it is common to observe heterogeneous variabilities between high-throughput samples, which violates the assumption of the Student’s *t*-test. For example, samples processed by a skillful technician usually have less variability than those processed by an inexperienced person. For two-group comparisons, a special case of the heterogeneity of variance, i.e., samples in different groups have different variances, is well studied and commonly referred to as the Behrens-Fisher problem. The best known (approximate) parametric solution for this problem is the Welch’s *t*-test, which adjusts the degrees of freedom (hereinafter, DFs) associated with the *t*-distribution to compensate for the heteroscedasticity in the data. Unfortunately, the Welch’s *t*-test is not appropriate when the data have even more complicated variance structure. As an example, it is well known that the quality and variation of the RNA-seq sample is largely affected by the total number of reads in the sequencing specimen (Wang et al., 2012; Sims et al., 2014). This quantity is also known as *sequencing depth* or *library size*, which may vary widely from sample to sample. Fortunately, such information is available *a priori* to data analyses. Several weighted methods (Law et al., 2014; Zhou et al., 2014; Liu et al., 2015) are proposed to utilize this information and make reliable statistical inference.

As the technology advances and the unit cost drops, immense amount of data are produced with even more complex variance-covariance structures. In multi-site studies for big data consortium projects, investigators sometimes need to integrate omics-data from different platforms (e.g. microarray or RNA-seq for gene expression) and/or processed in different batches. Although many normalization (Robinson & Oshlack, 2010; Risso et al., 2014; Liu et al., 2017) and batch-correction methods (Johnson et al., 2007; Leek & Storey, 2007; Hardcastle & Kelly, 2010) can be used to remove spurious *bias*, the heterogeneity of variance remains to be an issue. Besides, the clustering nature of these data may induce *correlation* among observations within one center/batch. Correlation may arise due to other reasons such as paired samples. For example, we downloaded a set of data for a comprehensive breast cancer study (Cancer Genome Atlas Network, 2012), which contain 226 samples including 153 tumor samples and 73 paired normal samples. Simple choices such as Welch’s *t*-test and paired *t*-test are not ideal for comparing the gene expression patterns between normal and cancerous samples, because they either ignore the correlations of the paired subjects or waste information contained in the unpaired subjects. To ignore the correlation and use a two-sample test imprudently is harmful because it may increase the type I error rate extensively (Walsh, 1947). On the other hand, a paired test can only be applied to the matched samples, which almost certainly reduces the detection power. In general, data that involves two or more matched samples are called repeated measurements, and it is very common in practice to have some unmatched samples, also known as unbalanced study design.

One of the most versatile tools in statistics, the linear mixed-effects regression (LMER), provides an alternative inferential framework that accounts both unequal variances and certain practical correlation structures. The standard LMER can model the correlation by means of random effects. By adding weights to the model, the weighted LMER is able to capture very complex covariance structures in real applications. Although LMER has many nice theoretical properties, fitting it is computationally intensive. Currently, the best implementation is the R package `lme4` (Bates et al., 2014), which is based on an iterative EM algorithm. For philosophical reasons, `lme4` does not provide *p*-values for the fitted models. The R package `lmerTest` (Kuznetsova et al., 2017) is the current practical standard to perform regression *t*- and *F*-tests for `lme4` outputs with appropriate DFs.

Many classical parametric tests, such as two-sample and paired *t*-tests, have their corresponding rank-based counterparts, i.e. the Wilcoxon rank-sum test and the Wilcoxon signed rank test. A rank-based solution to the Behrens-Fisher problem can be derived based on the adaptive rank approach (Sidak et al., 1999), but it was not designed for correlated observations. In recent years, researchers also extended rank-based tests to situations where both correlations and weights are presented. Barry et al. (2008) derived the Wilcoxon rank-sum statistic for correlated ranks, and Zhang et al. (2017) derived the weighted Mann-Withney U statistic for correlated data. These methods incorporate an interchangeable correlation in the whole dataset, and are less flexible for a combination of correlated and uncorrelated ranks. Lumley & Scott (2013) proved the asymptotic properties for a class of weighted ranks under complex sampling, and pointed out that a reference *t*-distribution is more appropriate than the normal approximation for the Wilcoxon test when the design has low DFs. Their method is implemented in the `svyranktest()` function in R package survey. But most of the rank-based tests are designed for group comparisons; rank-based approaches for testing associations between two continuous variables with complex covariance structure are underdeveloped.

Based on a linear regression model, we propose two H-T procedures (one parametric and one semiparametric) that utilize *a priori* information of the variance (weights) and correlation structure of the data. In Section 2, we design a linear map, dubbed as the “PB-transformation”, that a) transforms the original data with unequal variances and correlation into certain equivalent data that are independent and identically distributed; b) maps the original regression-like H-T problem into an equivalent *one-group* testing problem. After the PB-transformation, classical parametric and rank-based tests with adjusted DFs are directly applicable. We also provide a moment estimator for the correlation coefficient for repeated measurements, which can be used to obtain an estimated covariance structure if it is not provided *a priori*. In Section 3, we investigate the performance of the proposed methods using extensive simulations based on normal and double exponential distributions. We show that our methods have tighter control of type I error and more statistical power than a number of competing methods. In Section 4, we apply the PB-transformed *t*-test to an RNA-seq data for breast cancer. Utilizing the information of the paired samples and sequencing depths, our method selects more cancer-specific genes and fewer falsely significant genes (i.e. genes specific to other diseases) than the major competing method based on weighted LMER.

Lastly, computational efficiency is an important assessment of modern statistical methods. Depending on the number of hypotheses to be tested, our method can perform about 200 to 300 times faster than the LMER approach in simulation studies and real data analyses. This efficiency makes our methods especially suitable for fast feature selection in high-throughput data analysis.

## 2 Methods

### 2.1 Model Framework

For clarity, we first present our main methodology development for a univariate regression problem. We will extend it to multiple regression problems in Section 2.7.

Consider the following regression-type H-T problem:

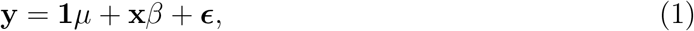

where *μ*, *β* ∈ ℝ, **y**, **x**, ***ϵ***, **1** = (1, · · ·, 1)′ ∈ ℝ^*n*^ and 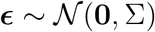

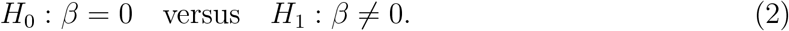

Here, **y** is the response variable, **x** is the covariate, and ***ϵ*** is the error term that follows an *n*-dimensional multivariate normal distribution 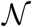 with mean zero and a general variance-covariance matrix Σ. By considering a random variable **Y** in the *n*-dimensional space, the above problem can also be stated as

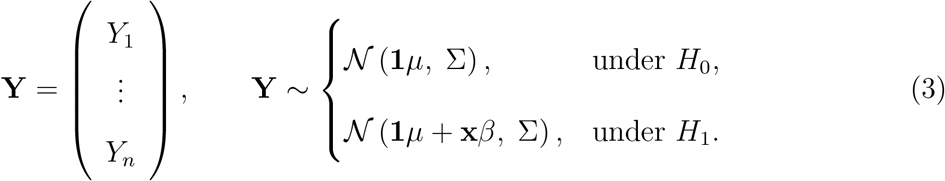

In this model, *μ* is the intercept or grand mean that is a nuisance parameter, and *β* is the parameter of interest that quantifies the effect size. We express the variance-covariance matrix of ***ϵ*** in the form

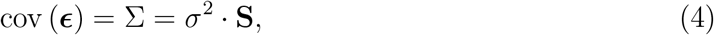

where *σ*^2^ is a nonzero scalar that quantifies the magnitude of the covariance structure, and **S** is a symmetric, positive-definite matrix that captures the *shape* of the covariance structure. Additional constraints are needed to determine *σ*^2^ and **S**; here, we choose a special form that can subsequently simplify our mathematical derivations. For any given Σ, define

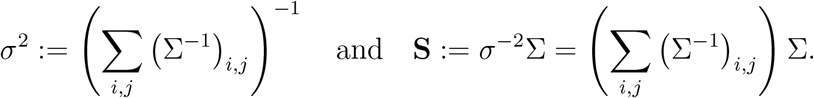

From the above definition, we have the following nice property

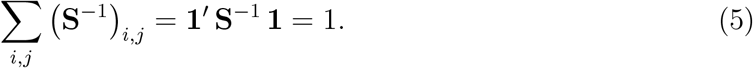

Hereinafter, we refer to **S** the *standardized* structure matrix satisfying Equation (5).

### 2.2 The Proposed Method

As a special case of Model (3), if **S** is proportional to **I**, the identity matrix, it is well-known that regression *t*-test is a valid solution to this H-T problem. If **S** ≠ **I**, e.g. the observed data are correlated and/or have heterogeneous variance structure, the assumptions of the standard *t*-test are violated. In this paper, we propose a linear transformation, namely **PB** : **Y** → **Ỹ**, which transforms the original data to a new set of data that are independent and identically distributed. Furthermore, we prove that the transformed H-T problem related to the new data is equivalent to the original problem, so that we can approach the original hypotheses using standard parametric (or later rank-based) tests with the new data.

To shed more lights on the proposed method, we first provide a graphical illustration in Figure 1. The proposed procedure consists of three steps.

1. Estimate 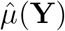 (i.e. the *weighted* mean of the original data), and subtract 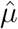 from all data. This process is an oblique (i.e. non-orthogonal) projection from ℝ^*n*^ to an (*n* − 1)-dimensional subspace of ℝ^*n*^. The intermediate data from this step is **Y**^(1)^ (i.e. the centered data). It’s clear that 𝔼**Y**^(1)^ is the origin of the reduced space if and only if *H*_0_ is true.
2. Use the eigen-decomposition of the covariance matrix of **Y** ^(1)^ to *reshape* its “elliptical” distribution to a “spherical” distribution. The intermediate data from this step is **Y**^(2)^.
3. Use the QR-decomposition technique to find a *unique rotation* that transforms the original H-T problem to an equivalent problem of testing for a constant deviation along the unit vector. The equivalent data generated from this step is **Ỹ**, and the H-T problem associated with **Ỹ**.can be approached by existing parametric and rank-based methods.

**Figure 1:**
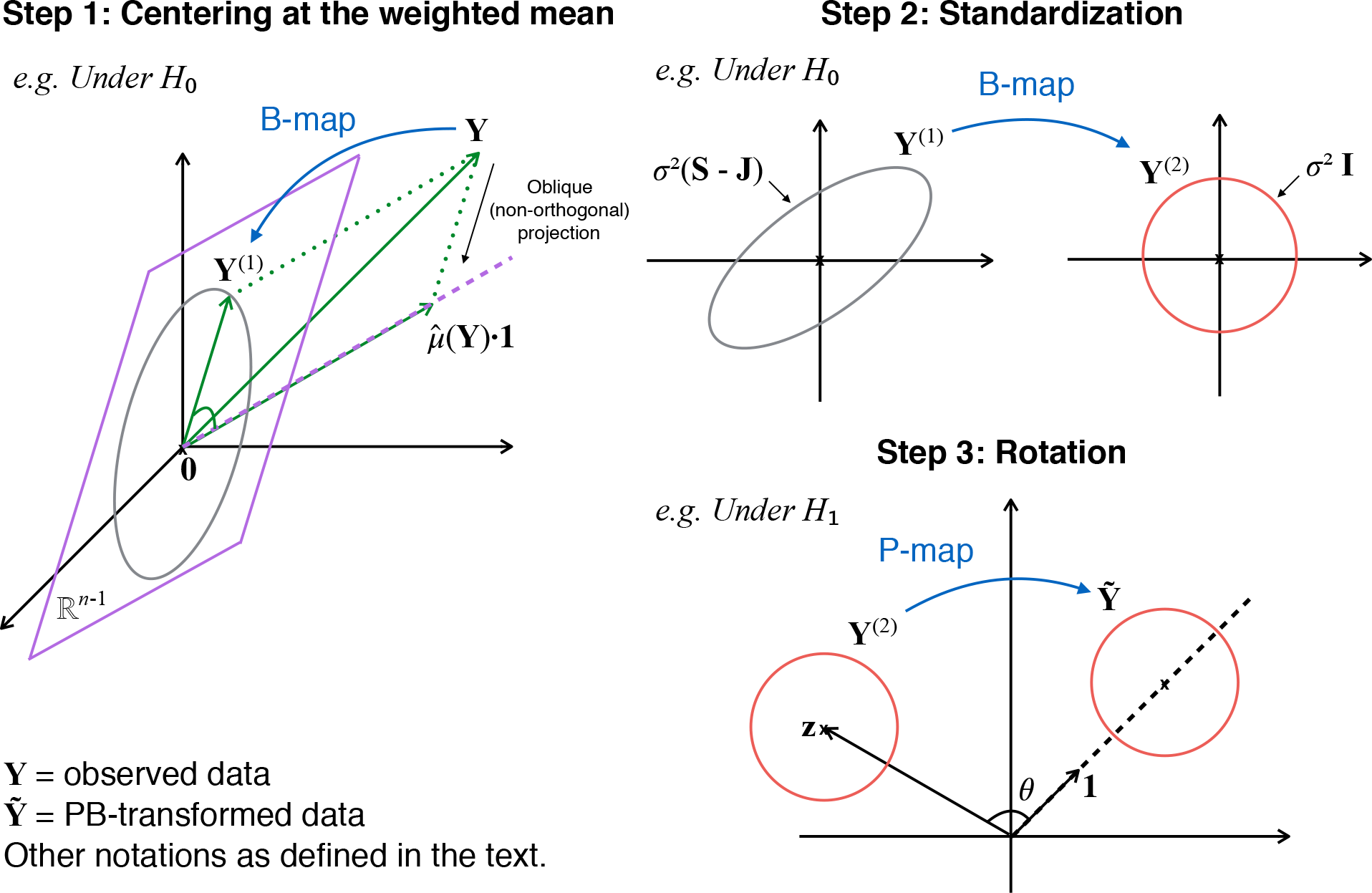
Graphical illustration of the PB-transformation. Step 1: Estimate 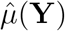 (i.e. the *weighted* mean of the original data), and subtract 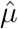 from all data. This process is an oblique (i.e. non-orthogonal) projection from R^*n*^ to an (*n* − 1)-dimensional subspace of R^*n*^. The intermediate data from this step is **Y**^(1)^, also call it centered data. If *H*_0_ is true, **Y**^(1)^ centers at the origin of the reduce space; otherwise, the data cloud **Y**^(1)^ deviates from the origin. Step 2: Use eigen-decomposition to *reshape* the “elliptical” distribution to an “spherical” distribution. The intermediate data from this step is **Y**^(2)^. Step 3: Use QR-decomposition to find a *unique rotation* that transforms the original H-T problem to an equivalent problem. The equivalent problem tests for a constant deviation along the unit vector in the reduced space, thus it can be approached by existing parametric and rank-based methods. The final data from this step is **Ỹ**.

In the proposed PB-transformation, B-map performs both transformations in Step 1 and 2; P-map from Step 3 is designed to improve the power of the proposed semiparametric test to be described in Section 2.6.

#### 2.2.1 Centering data

Using generalized least squares, the weighted mean for the original data is 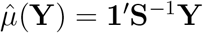. We subtract 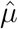 from all data points and define the centered data as

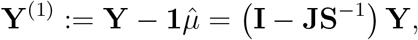

where **J** = **1** · **1′** (i.e. a matrix of all 1’s). With some mathematical derivations (see Supplementary Text, Section S1.1), we have

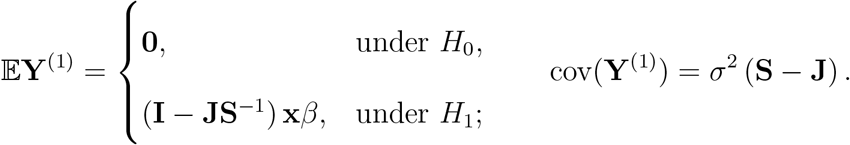

#### 2.2.2 The B-map

Now, we focus on **S** − **J**, which is the structure matrix of the centered data. Let **T** Λ**T′** denote the eigen-decomposition of **S** − **J**. Since the data are centered, there are only *n* − 1 nonzero eigenvalues. We express the decomposition as follows

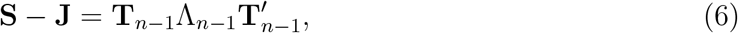

where **T**_*n*−1_ ∈ M_n×(n−1)_ is a semi-orthogonal matrix containing the first *n* − 1 eigenvectors and Λ_*n*−1_ ∈ M_(*n*−1)×(*n*−1)_ is a diagonal matrix of nonzero eigenvalues. Based on Equation (6),we define (see Supplementary Text, Section S1.2)

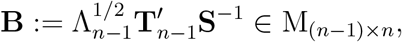

so that **Y**^(2)^ := **BY** ∈ ℝ^*n*−1^ have the following mean and covariance

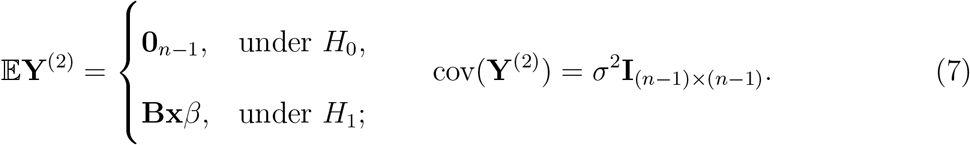

We call the linear transformation represented by matrix **B** the “B-map”. So far, we have centered the response variable, and standardized the general structure matrix **S** into the identity matrix **I**. However, the covariate and the alternative hypothesis in the original problem are also transformed by the B-map. For normally distributed **Y**, the transformed H-T problem in Equation (7) is approachable by the regression *t*-test; however, there’s no appropriate rank-based counterpart. In order to conduct a rank-based test for **Y** with broader types of distribution, we propose the next transformation.

#### 2.2.3 The P-map

From Equation (7), define the transformed covariate

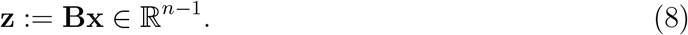

We aim to find an orthogonal transformation that aligns **z** to **1** _*n*−1_ in the reduced space. We construct such a transformation through the QR decomposition of the following object

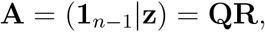

where **A** ∈ M_(*n*−1)×2_ is a column-wise concatenation of vector **z** and the target vector **1**_*n*−1_, **Q** ∈ M_(*n*−1)×2_ is a semi-orthogonal matrix, and **R** ∈ M_2×2_ is an upper triangular matrix. We also define the following rotation matrix

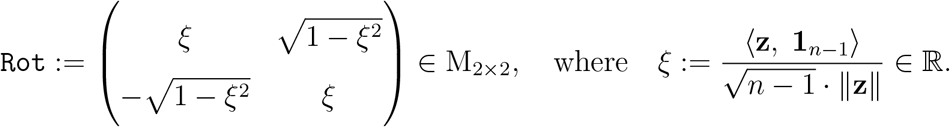

Geometrically speaking, *ξ* = cos *θ*, where *θ* is the angle between **z** and **1**_*n*−1_.

With the above preparations, we have the following result.

##### Theorem 2.1

*Matrix* **P** := **I** − **QQ**′ + **Q** `Rot` **Q**′ = **I** _(*n*−1)×(*n*−1)_ − **Q** (**I** _2×2_ − `Rot`)**Q**′ *is the unique orthogonal transformation that satisfies the following properties:*

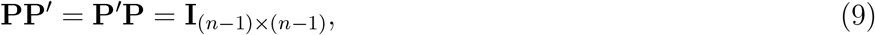

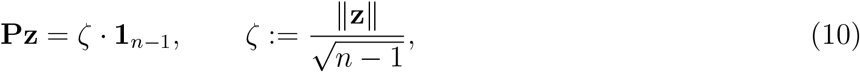

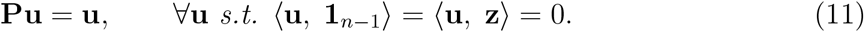

*Proof.* See Supplementary Text, Section S1.3.

We call the linear transformation **P** defined by Theorem 2.1 the “P-map”. Equation (9) ensures that this map is an orthogonal transformation. Equation (10) shows that the vector **z** is mapped to **1**_*n*−1_ scaled by a factor *ζ*. Equation (11) is an invariant property in the linear subspace 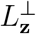, which is the orthogonal complement of the linear subspace spanned by **1**_*n*−1_ and **z**, i.e. *L*_**z**_ = span(**1**_*n*−1_, **z**). This property defines a *unique minimum* map that only transforms the components of data in *L*_**z**_ and leaves the components in 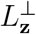 invariant.

With both **B** and **P**, we define the final transformed data as **Ỹ** := **PY**^(2)^ = **PBY**, which has the following joint distribution

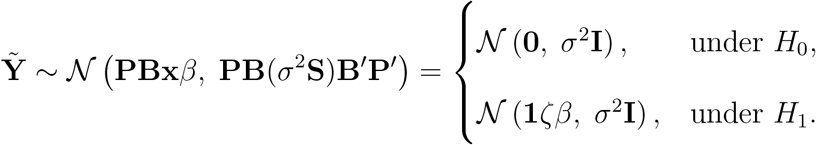

The normality assumption implies that each *Ỹ*_*i*_ follows an *i.i.d.* normal distribution, for *i* = 1, · · ·, *n* − 1. The location parameter of the common marginal distribution is to be tested with unknown *σ*^2^. Therefore, we can approach this equivalent H-T problem with the classical one-sample *t*-test and Wilcoxon signed rank test (more in Section 2.6).

### 2.3 Correlation estimation for repeated measurements

If Σ is unknown, we can decompose Σ in the following way

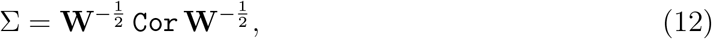

where **W** is a diagonal weight matrix and `Cor` is the corresponding correlation matrix. By definition, the weights are inversely proportional to the variance of the observations. In many real world applications including RNA-seq analysis, those weights can be assigned *a priori* based on the quality of samples; but the correlation matrix `Cor` needs to be estimated from the data. In this section, we provide a moment-based estimator of `Cor` for a class of correlation structure that is commonly used for repeated measurements. This estimator does not require computationally intensive iterative algorithms.

Let **Y** be a collection of repeated measures from *L* subjects such that the observations from different subjects are independent. With an appropriate data rearrangement, the correlation matrix of **Y** can be written as a block-diagonal matrix

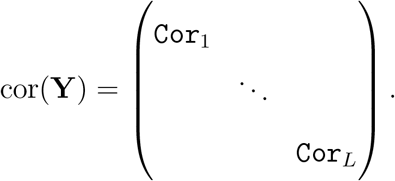

We assume that the magnitude of correlation is the same across all blocks, and denote it by *ρ*. Each block can be expressed as 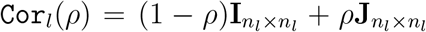 for *l* = 1*, · · ·, L,* where *n*_*l*_ is the size of the *l*th block and 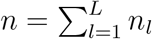

We estimate the correlation based on the weighted regression residuals 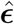 defined by Equation (S3) in Supplementary Text, Section S2.1. Define two forms of residual sum of squares

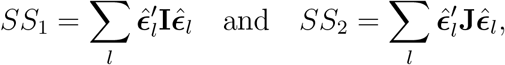

where 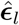 is the corresponding weighted residuals for the *l*th block. With these notations, we have the following Proposition.

#### Proposition 2.2

*Denote* 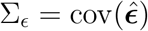 *and assume that for some nonzero σ*^2^,

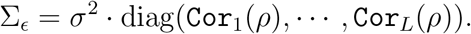

*An estimator of ρ based on the first moments of SS*_1_ *and SS*_2_ *is*

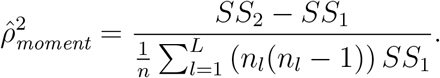

*Moreover, if* 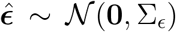 *and n*_1_ = · · · = *n*_*L*_ = *n/L (i.e. balanced design), the above estimator coincides with the maximum likelihood estimator of ρ, which has the form*

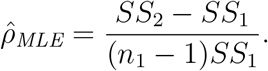

*Proof.* See Supplementary Text, Section S2.1.

Standard correlation estimates are known to have downward bias (Zimmerman et al., 2003), which can be corrected by the Olkin and Pratt’s method (Olkin & Pratt, 1958). With this correction, our final correlation estimator is

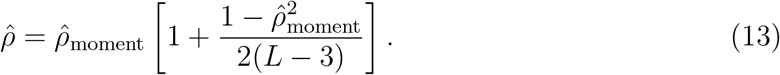

### 2.4 Kenward-Roger approximation to the degrees of freedom

The degree of freedom (DF) can have nontrivial impact on hypothesis testing when sample size is relatively small. Intuitively, a correlated observation carries “less information” than that of an independent observation. In such case, the effective DF is smaller than the apparent sample size. Simple examples include the two-sample *t*-test and the paired *t*-test. Suppose there are *n* observations in each group, the former test has DF = 2*n* − 2 for *i.i.d.* observations, and the latter only has DF = *n* − 1 because the observations are perfectly paired. These trivial examples indicate that we need to adjust the DF according to the correlation structure in our testing procedures.

We adopt the degrees of freedom approximation proposed by Kenward & Roger (1997) (K-R approximation henceforth) for the proposed tests. The K-R approximation is a fast moment-matching method, which is efficiently implemented in R package pbkrtest (Halekoh et al., 2014). In broad terms, we use the DF approximation as a tool to adjust the effective sample size when partially paired data are observed.

### 2.5 Alternative approach using mixed-effects model

As we mentioned in Section 1, the H-T problem stated in Model (3) for repeated measurements can also be approached by the linear mixed-effects regression (LMER) model. Suppose the *i*th observation is from the *l*th subject, we may fit the data with a random intercept model such that

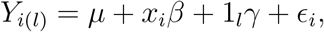

where 1_*l*_ is the indicator function of the *l*th subject, 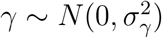, and 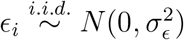. The correlation is modeled as

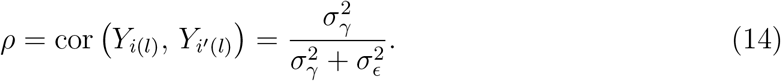

The LMER model is typically fitted by a likelihood approach based on the EM algorithm. Weights can be incorporated in the likelihood function. The `lmer()` function in R package `lme4` (Bates et al., 2014) provides a reference implementation for fitting the LMER model. The algorithm is an iterative procedure until convergence. Due to relatively high computational cost, the mixed-effects model has limited application in high-throughput data.

The R package `lmerTest` (Kuznetsova et al., 2017) performs hypothesis tests for `lmer()` outputs. By default, it adjusts the DF using the Satterthwaites approximation (Satterthwaite, 1941), and can optionally use the K-R approximation.

### 2.6 A Semiparametric Generalization

In the above sections, we develop the PB-transformed *t*-test using linear algebra techniques. These techniques can be applied to non-normal distributions to transform their mean vectors and covariance matrices as well. With the following proposition, we may extend the proposed method to an appropriate semiparametric distribution family. By considering the uncorrelated observations with equal variance as a second order approximation of the data that we are approaching, we can apply a rank-based test on the transformed data to test the original hypotheses. We call this procedure the PB-transformed Wilcoxon test.

#### Proposition 2.3

*Let* 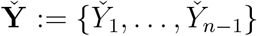 *be a collection of i.i.d. random variables with a common symmetric density function g*(*y*), *g*(−*y*) = *g*(*y*)*. Assume that* 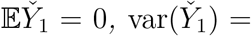, 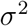. *Let Y*\* *be a random number that is independent of* 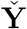 *and has zero mean and variance σ*^2^. *For every symmetric semi-definite* **S** *∈* M_*n×n*_, **x** *∈* ℝ^*n*^ *and μ, β ∈* ℝ, *there exists a linear transformation* **D** : ℝ^*n−*1^ → ℝ^*n*^ *and constants u, v, such that*

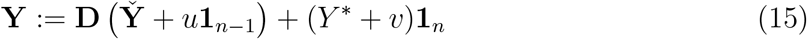

*is an n-dimensional random vector with*

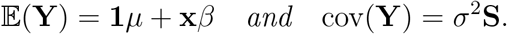

*Furthermore, if we apply the PB-transformation to* **Y** *, the result is a sequence of* (*n* − 1) *equal variance and uncorrelated random variables with zero mean if and only if β* = 0. *Proof.*

See Supplementary Text, Section S1.4.

The essence of this Proposition is that, starting with an *i.i.d.* sequence of random variables with a symmetric common p.d.f., we can use linear transformations to generate a family of distributions that is expressive enough to include a non-normal distribution with an arbitrary covariance matrix and a mean vector specified by the effect to be tested. This distribution family is *semiparametric* because: a) the “shape” of the density function, *g*(*y*), has infinite degrees of freedom; b) the “transformation” (**D**, *u*, and *v*) has only finite parameters.

As mentioned before, applying both the B- and P-maps enables us to use the Wilcoxon signed rank test for the hypotheses with this semiparametric distribution family. This approach has better power than the test with only the B-map as shown in Section 3. Once the PB-transformed data are obtained, we calculate the Wilcoxon singed rank statistic and follow the testing approach in Lumley & Scott (2013), which is to approximate the asymptotic distribution of the test statistic by a *t*-distribution with an adjusted DF. In summary, this PB-transformed Wilcoxon test provides an approximate test (up to the second order moment) for data that follow a flexible semiparametric distributional model.

### 2.7 Extension to Multiple Regressions

In this section, we present an extension of the proposed methods for the following multiple regression

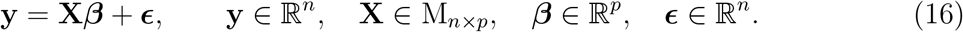

Here the error term *ϵ* is assumed to have zero mean but does not need to have scalar covariance matrix. For example, *ϵ* can be the summation of random effects and measurement errors in a typical LMER model with a form specified in Equation (4).

To test the significance of *β*_*k*_, *k* = 1, …, *p*, we need to specify two regression models, the null and alternative models. Here the alternative model is just the full Model (16), and the null model is a regression model for which the covariate matrix is **X**_−*k*_, which is constructed by removing the *k*th covariate (*X*_*k*_) from **X**

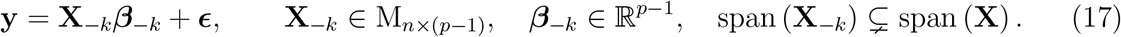

Compared with the original univariate problem, we see that the *nuisance covariates* in the multiple regression case are **X**_−*k*_**β**_−*k*_ instead of **1***μ* in Equation (1). Consequently, we need to replace the centering step by regressing out the linear effects of **X**_−*k*_

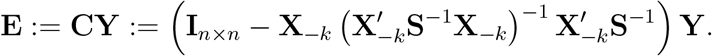

The new B-transformation is defined as the eigen-decomposition of cov 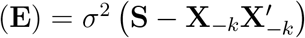

The P-transformation is derived the same as before, but with the new **B** matrix.

## 3 Simulations

We design two simulation scenarios for this study: SIM1 for completely paired group comparison, and SIM2 for regression-type test with a continuous covariate. For both scenarios we consider two underlying distributions (normal and double exponential) and two correlation levels (*ρ* = 0.2 and *ρ* = 0.8). We compare the parametric and rank-based PB-transformed test with oracle and estimated correlation to an incomplete survey of alternative methods. Each scenario was repeated ten times and the results are summarized in Figures 2 and 3, and Tables 1 and 2. See Supplementary Text, Section S3 for more details.

**Table 1:** Type-I error and power comparison for group comparison tests. At the 5% significance level, mean and standard deviation (in brackets) of the type-I error rate and power over 10 sets of SIM1 data are reported.

| *SIM1: Group Comparison* | $\rho = 0.2$ | | $\rho = 0.8$ | |
| --- | --- | --- | --- | --- |
|  | Type-I error | Power | Type-I error | Power |
| **Normal** |  |  |  |  |
| PB.t (oracle) | 0.037 (0.005) | 0.844 (0.012) | 0.035 (0.005) | 0.922 (0.008) |
| PB.t (estimation) | 0.037 (0.005) | 0.843 (0.013) | 0.028 (0.005) | 0.913 (0.010) |
| Weighted LMER | 0.042 (0.006) | 0.829 (0.014) | 0.022 (0.003) | 0.613 (0.013) |
| Weighted regression $t$-test | 0.032 (0.004) | 0.824 (0.014) | 0.002 (0.001) | 0.365 (0.020) |
| Paired $t$-test | 0.050 (0.007) | 0.502 (0.014) | 0.048 (0.003) | 0.468 (0.012) |
| Two-sample $t$-test | 0.032 (0.005) | 0.456 (0.011) | 0.000 (0.001) | 0.092 (0.009) |
| **Double exponential** |  |  |  |  |
| PB.wilcox (oracle) | 0.034 (0.006) | 0.896 (0.011) | 0.032 (0.006) | 0.949 (0.010) |
| PB.wilcox (estimation) | 0.047 (0.009) | 0.862 (0.013) | 0.034 (0.006) | 0.915 (0.010) |
| svyranktest | 0.116 (0.009) | 0.820 (0.009) | 0.099 (0.009) | 0.616 (0.014) |
| Friedman | 0.052 (0.006) | 0.492 (0.016) | 0.053 (0.006) | 0.518 (0.016) |
| Wilcoxon signed rank | 0.055 (0.007) | 0.567 (0.017) | 0.058 (0.009) | 0.566 (0.012) |
| Wilcoxon rank-sum | 0.039 (0.007) | 0.600 (0.015) | 0.002 (0.002) | 0.216 (0.013) |

**Table 2:** Type-I error and power comparison for regression tests. At the 5% significance level, mean and standard deviation (in brackets) of the type-I error rate and power over 10 sets of SIM2 data are reported.

| *SIM2: Regression Test* | $\rho = 0.2$ | | $\rho = 0.8$ | |
| --- | --- | --- | --- | --- |
|  | Type-I error | Power | Type-I error | Power |
| **Normal** |  |  |  |  |
| PB.t (oracle) | 0.044 (0.007) | 0.756 (0.015) | 0.042 (0.007) | 0.692 (0.018) |
| PB.t (estimation) | 0.044 (0.007) | 0.756 (0.016) | 0.036 (0.006) | 0.671 (0.018) |
| Weighted LMER | 0.050 (0.009) | 0.751 (0.019) | 0.051 (0.006) | 0.603 (0.016) |
| Weighted regression *t*-test | 0.049 (0.008) | 0.750 (0.016) | 0.054 (0.007) | 0.400 (0.016) |
| Regression *t*-test | 0.051 (0.007) | 0.684 (0.015) | 0.051 (0.007) | 0.356 (0.019) |
| **Double exponential** |  |  |  |  |
| PB.wilcox (oracle) | 0.040 (0.008) | 0.817 (0.011) | 0.037 (0.007) | 0.736 (0.013) |
| PB.wilcox (estimation) | 0.060 (0.009) | 0.727 (0.018) | 0.068 (0.009) | 0.623 (0.016) |
| B.spearman (estimation) | 0.079 (0.011) | 0.678 (0.020) | 0.079 (0.011) | 0.588 (0.022) |
| Spearman test | 0.073 (0.010) | 0.641 (0.017) | 0.072 (0.009) | 0.328 (0.014) |

**Figure 2:**
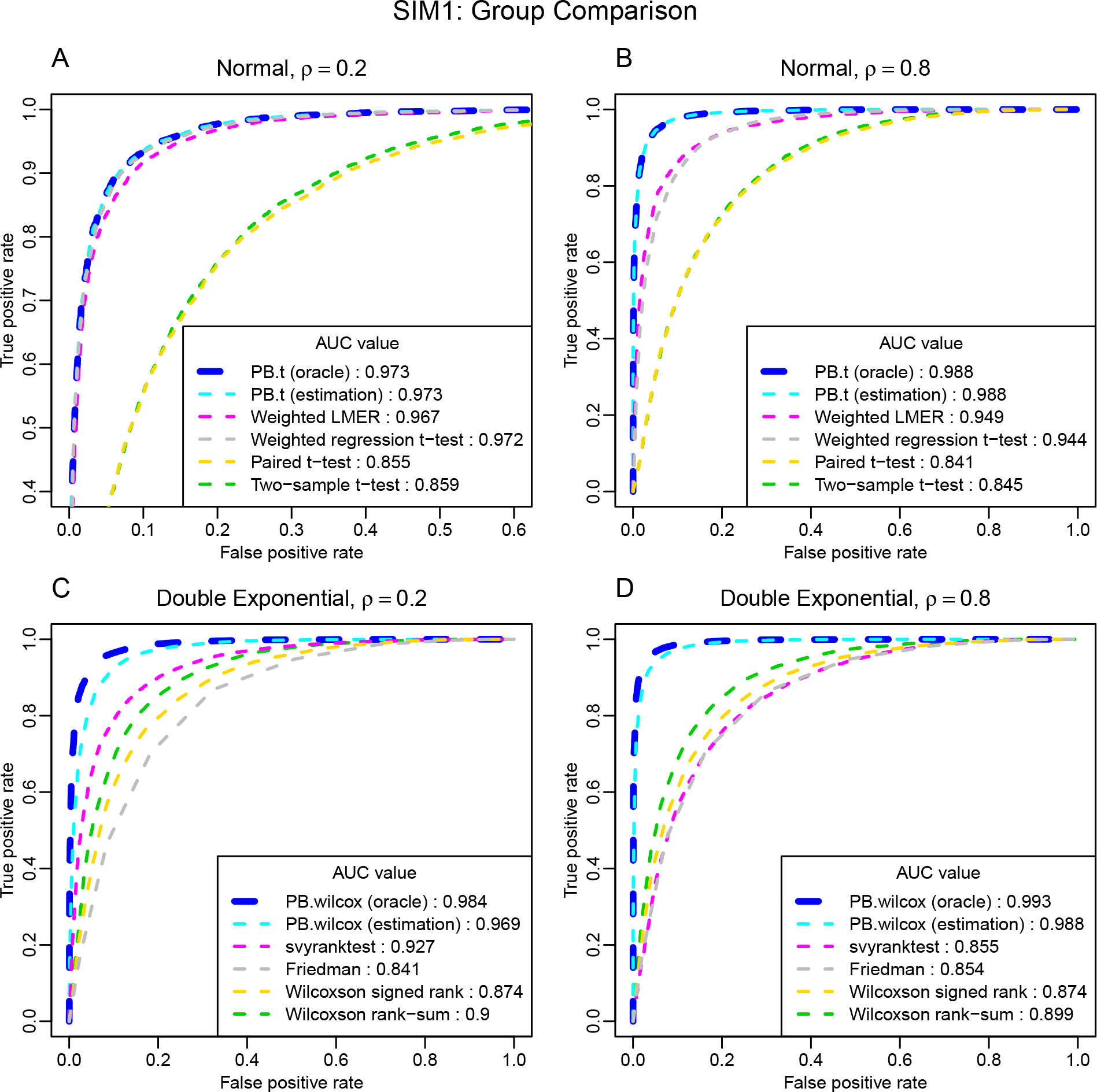
ROC curves for group comparison tests. In SIM1, six parametric methods and six rank-based methods are compared. AUC values are reported in the legend. Plot A is zoomed to facilitate the view of curves that overlay on top of each other. For both *ρ* = 0.2 and *ρ* = 0.8, the PB-transformed parametric and rank-based tests outperform all other tests.

**Figure 3:**
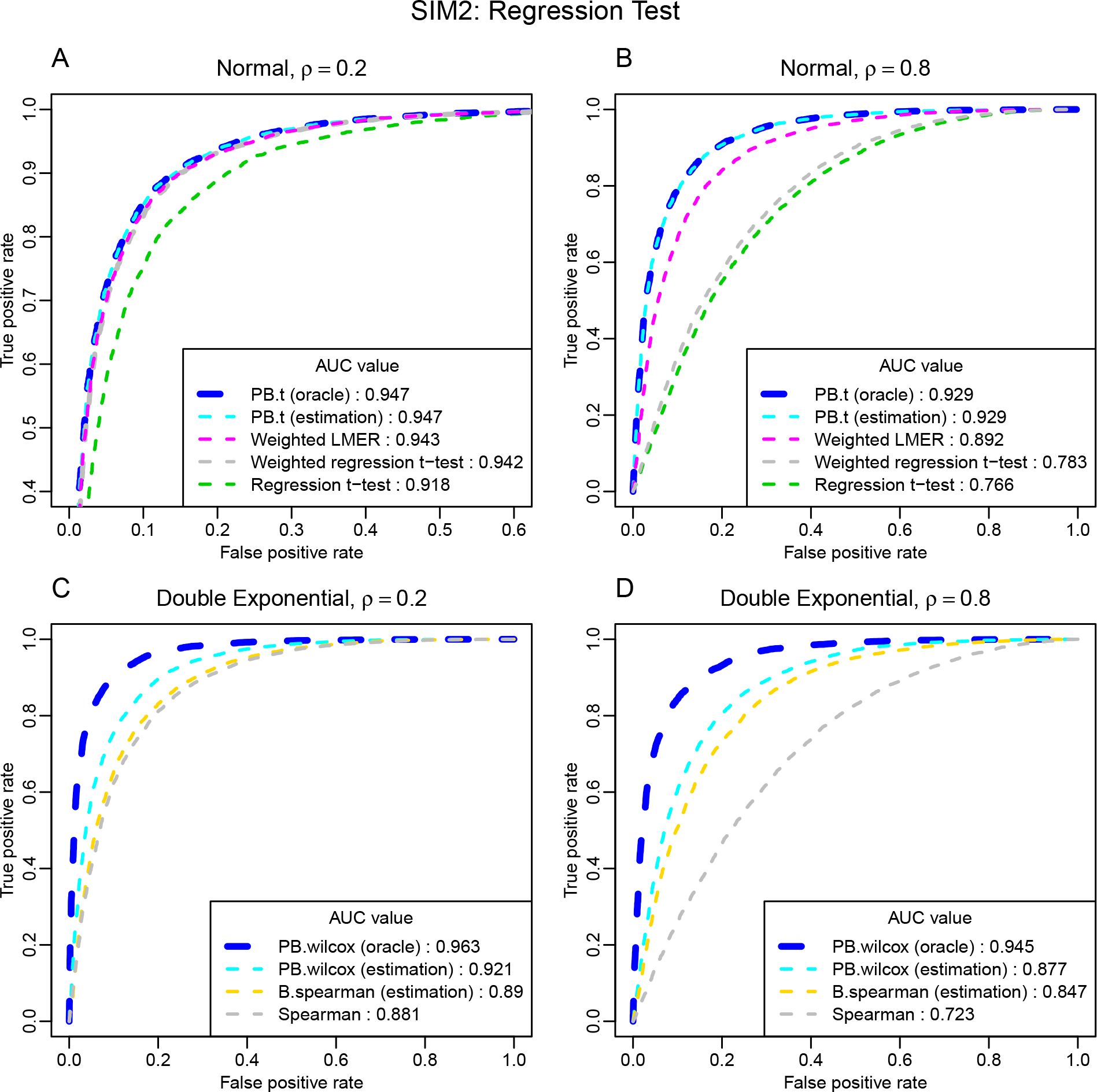
ROC curves for regression tests. In SIM2, five parametric methods and four rank-based methods are compared. AUC values are reported in the legend. Plot A is zoomed to facilitate the view of curves that overlay on top of each other. For both *ρ* = 0.2 and *ρ* = 0.8, the PB-transformed parametric and rank-based tests outperform all other tests.

Figure 2 and Figure 3 are ROC curves for SIM1 and SIM2, respectively. In all simulations, the proposed PB-transformed tests outperform the competing methods.

The PB-transformed *t*-test has almost identical performance with oracle or estimated *ρ*. Using the estimated *ρ* slightly lowers the ROC curve of the PB-transformed Wilcoxon test compared with the oracle curve, but it still has a large advantage over other tests. Within the parametric framework, the weighted LMER has the best performance among the competing methods. It achieves similar performance as our proposed parametric test when the correlation coefficient is small; however, its performance deteriorates when the correlation is large. Judging from the ROC curves, among the competing methods, the `svyranktest()` is the best rank-based test for the group comparison problem, primarily because it is capable of incorporating the correlation information. However, it fails to control the type-I error, as shown in Table 1.

Table 1 and Table 2 summarize the type-I error rate and power at the 5% significance level for SIM1 and SIM2, respectively. Overall, the PB-transformed tests achieve the highest power in all simulations. In most cases, the proposed tests tend to be conservative in the control of type-I error; and replacing the oracle *ρ* by the estimated 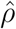 does not have significant impact on the performance of PB-transformed tests. The only caveat is the rank-based test for the regression-like problem. Currently, there’s no appropriate method designed for this type of problem. When the oracle correlation coefficient is provided to the PB-transformed Wilcoxon test, it has tight control of type I error. With uncertainty in the estimated correlation coefficient, our PB-transformed Wilcoxon test may suffer from slightly inflated type I errors; but it is still more conservative than its competitors. Of note, other solutions, such as the naive *t*-test and rank-based tests, may have little or no power for correlated data, though they may not have the lowest ROC curve.

### 3.1 Computational cost and degrees of freedom

We record the system time for testing 2,000 simulated hypotheses using our method and `lmer()`. Our method takes less than 0.3 second with given Σ, and less than 0.9 second with the estimation step; `lmer()` takes 182 seconds. We use a MacBook Pro equipped with 2.3 GHz Intel Core i7 processor and 8GB RAM (R platform: x86 64-darwin15.6.0). Of note, `lmer()` may fail to converge occasionally, e.g. 0 – 25 failures (out of 2,000) in each repetition of our simulations. We resort to a `try/catch` structure in the R script to prevent these convergence issues from terminating the main loop.

We also check the degrees of freedom in all applicable tests. In this section, we report the DFs used/adjusted in SIM1, i.e. the completely paired group comparison. Recall that *n* = 40 with *n*_A_= *n*_B_= 20. It is straightforward to calculate the DFs used in the two-sample *t*-test and the paired *t*-test, which are 38 and 19, respectively. Using `lmerTest()` (weighted LMER) with default parameters, it returns the mean DF = 35.51 with a large range (min = 4.77, max = 38) from the simulated data with *ρ* = 0.2. Using the oracle Σ_SIM_, our method returns the adjusted DF = 14.35; if the covariance matrix is estimated, our method returns the mean DF = 14.38 with high consistency (min = 14.36, max = 14.42). When *ρ* = 0.8, the adjusted DFs become smaller. The weighted LMER returns the mean DF = 20.63 (min = 4.03, max = 38). Our method returns DF = 12.48 for the oracle covariance, and mean DF = 12.56 (min = 12.55, max = 12.57) for the estimated covariance. Also, the rank-based test `svyranktest()` returns a DF for its *t*-distribution approximation, which is 18 for both small and large correlations.

## 4 A Real Data Application

We download a set of RNA-seq gene expression data from The Cancer Genome Atlas (TCGA) (Cancer Genome Atlas Network, 2012). The data are sequenced on the Illumina GA platform with tissues collected from breast cancer subjects. In particular, we select 28 samples from the tissue source site “BH”, which are controlled for white female subjects with the HER2-positive (HER2+) (Burstein, 2005) biomarkers. After data preprocessing based on nonspecific filtering, a total number of 11,453 genes are kept for subsequent analyses. Among these data are 10 pairs of matched tumor and normal samples, 6 unmatched tumor samples, and 2 unmatched normal samples. Using Equation (13), the estimated correlation between matched samples across all genes is 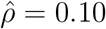.

The sequencing depths of the selected samples range from 23.80 million reads to 76.08 million reads. As mentioned before, the more reads are sequenced, the better is the quality of RNA-seq data (Sims et al., 2014); thus it is reasonable to weigh samples by their sequencing depths. Since this quantity is typically measured in million reads, we set the weights

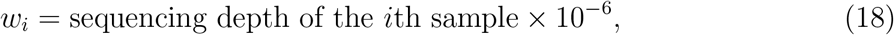

for *i* = 1, · · ·, 28.

With the above correlation estimate and weights, we obtained the covariance structure using Equation (12). For properly preprocessed sequencing data, a proximity of normality can be warranted (Ritchie et al., 2015). We applied the PB-transformed *t*-test and the weighted LMER on the data.

Based on the simulations, we expect that if correlation is small, the PB-transformed *t*-test should have tighter control of false positives than alternative methods. At 5% false discovery rate (FDR) level combined with a fold-change (FC) criterion (FC < 0.5 or FC > 2), the PB-transformed *t*-test selected 3,340 DEGs and the weighted LMER selected 3,485 DEGs.

To make the comparison between these two methods more fair and meaningful, we focus on studying the biological annotations of the top 2,000 genes from each DEG list. Specifically, we apply the gene set analysis tool DAVID (Huang et al., 2009a) to the 147 genes that uniquely belong to one list. Both Gene Ontology (GO) biological processes (Ashburner et al., 2000) and KEGG pathways (Kanehisa & Goto, 2000) are used for functional annotations. Terms identified based on the 147 unique genes in each DEG list are recorded in Supplementary Table S2. We further pin down two gene lists, which consist of genes that participate in more than five annotation terms in the above table: there are 11 such genes (PIK3R2, AKT3, MAPK13, PDGFRA, ADCY3, SHC2, CXCL12, CXCR4, GAB2, GAS6, and MYL9) for the PB-transformed *t*-test, and six (COX6B1, HSPA5, COX4I2, COX5A, UQCR10, and ERN1) for the weighted LMER. Expression level of these genes are plotted in Figure 4. These DEGs are biologically important because they are involved in multiple biological pathways/ontology terms.

**Figure 4:**
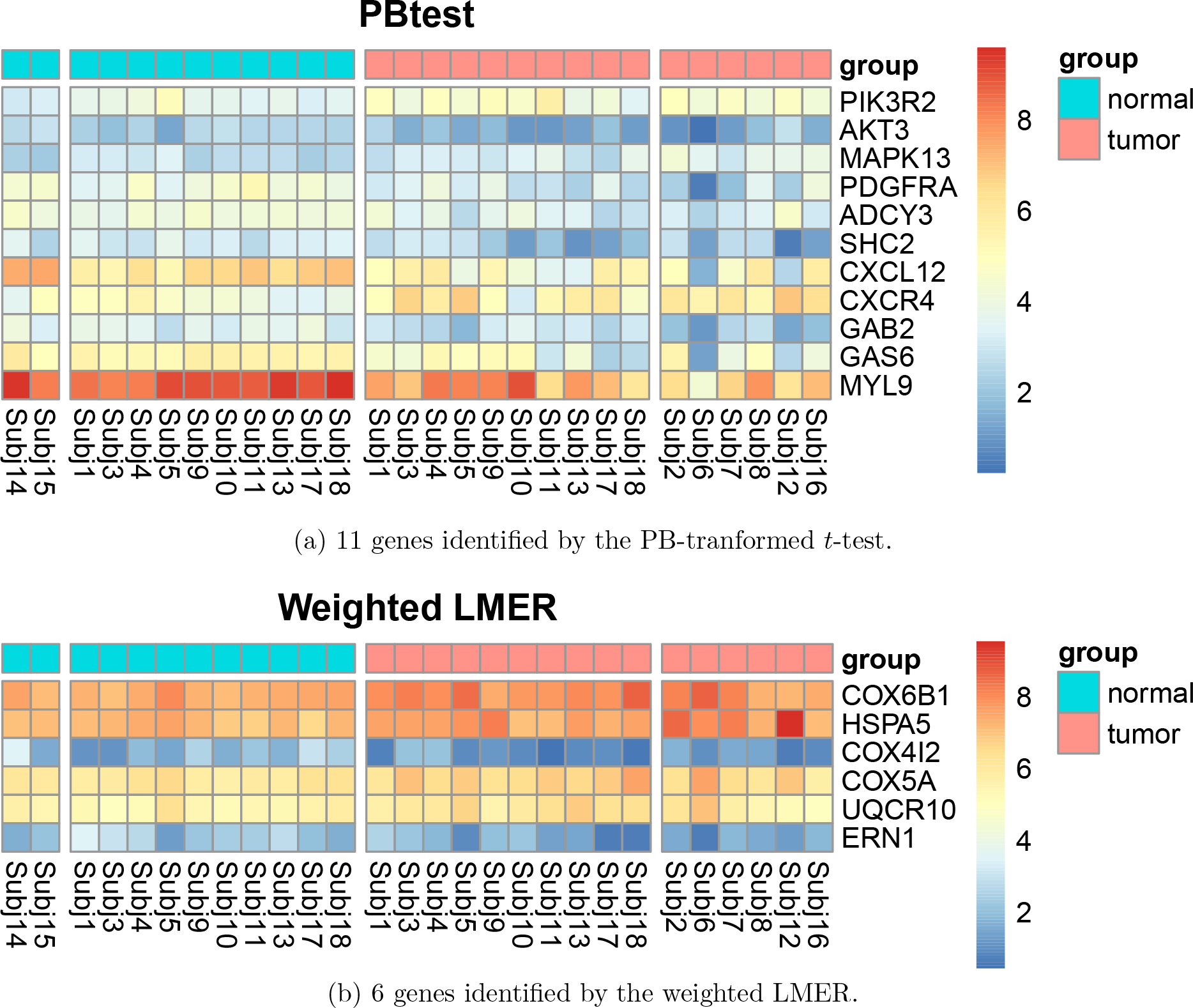
Selected differentially expressed genes uniquely identified by each test. Genes are in rows, and samples are in columns. The columns are ordered as unmatched normal samples, matched normal samples, matched tumor samples, and unmatched tumor samples. The selected genes are those who participated in more than five functional annotations in Supplementary Table S2. These genes are not only differentially expressed, but also biologically meaningful.

Those 11 genes uniquely identified by the PB-transformed *t*-test are known to be involved in cell survival, proliferation and migration. The CXCR4-CXCL12 chemokine signaling pathway is one of the deregulated signaling pathway uniquely identified by PB-transformed *t*-test in HER2+ breast cancer cells. This pathway is known to play a crucial role in promoting breast cancer metastasis and has been reported to be associated with poor prognosis (Sun et al., 2014; Müller et al., 2001). Compared with the state-of-the-art method (weighted LMER), the PB-transformed *t*-test identifies more genes whose protein products can be targeted by pharmaceutical inhibitors. CXCR4 inhibitors have already demonstrated promising anti-tumor activities against breast (Huang et al., 2009b; Chittasupho et al., 2017), prostrate (Wong et al., 2014) and lung (Taromi et al., 2016) cancers. Additional downstream signaling molecules identified by our analysis to be significantly associated with HER2+ breast tumor such as PI3K, p38, adaptor molecule GAB2 and SHC2 can also be potential therapeutic targets for selectively eliminating cancer cells. Please refer to Supplementary Text, Section S4.4 for more detailed biological explanations of these DEGs.

## 5 Discussion

In this paper, we present a data transformation technique that can be used in conjunction with both the Student’s *t*-type test and rank-based test. In the simulation studies, our proposed tests outperform the classical tests (e.g. two-sample/regreesion *t*-test and Wilcoxon rank-sum test) by a large margin. In a sense, this superiority is expected, because the classical methods do not consider the correlation nor heteroscedasticity of the data.

In our opinion, the most practical comparison in this study is the one between the PB-transformed *t*-test and the weighted LMER. The fact that the PB-transformed *t*-test outperforms the weighted LMER, and this advantage is more pronounced for data with higher correlation (see e.g., Figure 2 and 3), is the highlight of this study, which may have profound implications for applied statistical practice.

We believe the following reasons may explain the advantages of the PB-transformed tests.

1. As reported in Section 3.1, the default degrees of freedom approximation in `lmerTest` varies dramatically, as oppose to very stable degrees of freedom approximation in our method.
2. Our moment-based correlation estimator is better than the LMER correlation estimator (see Supplementary Text, Section S2.2). One possible explanation is that LMER depends on nonlinear optimizer, which may not always converge to the *global* maximum likelihood.
3. In a minor way but related to 2, `lmer()` fails to converge to even a *local* maximum in certain rare cases.

Another major contribution of our method is that the transformation-based approach is computationally much more efficient than the EM algorithm used in LMER, which is an important advantage in high-throughput data analysis. Recall that in simulation studies, PB-transformed *t*-test is approximately 200 times faster than the weighted LMER approach. As an additional evidence, to test the 11,453 genes in the real data study, it takes 933 seconds using the weighted LMER, and only 3 seconds using our method, which is more than 300 times faster.

Nonetheless, we want to emphasize that, by no means, our method is a replacement for LMER. The mixed-effects model is a comprehensive statistical inference framework that includes parameter estimation, model fitting (and possibly model selection), hypothesis testing, among other things; whereas our methods are only designed for the hypothesis testing. We envision that in a typical high-throughput data application, an investigator may quickly run PB-transformed *t*-test to identify important features first, then apply `lme4` to fit mixed effects models for those selected features. In this way, he/she enjoys both the computational efficiency of our method and the comprehensive results provided by a full LMER model.

In Section 2.7, we extend the PB-transformed tests for multiple regressions. We must point out two weaknesses in this approach. 1. The proposed extension is comparable to the regression *t*-test for individual covariates, not the ANOVA *F*-test for the significance of *several* covariates simultaneously. In fact, the B-map can be defined in this case so we can define a transformed parametric test easily; but there is no clear counterpart for the *P*-map, which is needed to overcome the identifiability issue for the semiparametric generalization. 2. The performance of PB-transformations depends on a good estimation of *S*, the shape of the covariance matrix of the observations. Currently, our moment-based estimator only works for problems with just one random intercept, which is only appropriate for relatively simple longitudinal experiments with limited (2 ~ 3) repeated measures. It is a challenging problem to estimate the complex covariance structure for general LMER models (e.g., one random intercept plus several random slopes), and we think it can be a nice and ambitious research project for us in the near future.

Numerically, the PB-transformed *t*-test provides the same test statistic and degrees of freedom as those from the paired *t*-test for perfectly paired data and the regression *t*-test for *i.i.d.* data. In this sense, the PB-transformed *t*-test is a legitimate generalization of these two classical tests. The rank-based test is slightly different from the classical ones, since we used a *t*-distribution approximation instead of a normal approximation for the rank-based statistic. The *t*-distribution approximation is preferred for correlated data because the *effective* sample size may be small even in a large dataset (Lumley & Scott, 2013).

Recall that the PB-transformation is designed in a way that the transformed data have the desired first and second order moments. For non-normal distributions, the transformed samples may not have the same higher order moments. Note that, the P-map is currently defined in part by Equation (11), the minimum action principle. Without this constraint, we will have some extra freedom in choosing the P-map. In the future development, we will consider using this extra freedom of orthogonal transformation to minimize the discrepancy of higher order moments of the transformed samples for the semiparametric distribution family. This would require an optimization procedure on a sub-manifold of the orthogonal group, which may be computationally expensive. The advantage is that, by making the higher order moments more homogeneous across the transformed data, we may be able to further improve the statistical performance of the PB-transformed Wilcoxon test.

In this study, we presented an example in RNA-seq data analysis. In recent bioinformatics research, advanced methods such as normalization and batch-effect correction were developed to deal with data heterogeneities in bio-assays. While most of these approaches are focused on the first moment (i.e. correction for bias in the mean values), our approach provides a different perspective based on the second order moments (i.e. the covariance structure). The dramatic computational efficiency boost of our method also opens the door for investigators to use the PB-transformed tests for ultra-high-dimensional data analysis, such as longitudinal studies of diffusion tensor imaging data at the voxel-level (Zhu et al., 2013; Liu et al., 2016a,b), in which about one million hypotheses need to be tested simultaneously. Finally, we think the PB-transformed Wilcoxon test can also be used in meta-analysis to combine results from several studies with high between-site variability and certain correlation structure due to, e.g., site- and subject-specific random effects.

## Acknowledgments

Research reported in this publication was supported in part by the National Institute of Environmental Health Sciences of the National Institutes of Health (NIH) under award number T32ES007271, the University of Rochester CTSA award number UL1 TR002001 from the National Center for Advancing Translational Sciences of the National Institutes of Health, the University of Rochester Center for AIDS Research (NIH 5 P30 AI078498-08), and Respiratory Pathogens Research Center (NIAID contract number HHSN272201200005C). The content is solely the responsibility of the authors and does not necessarily represent the official views of the NIH.

## Supplementary Materials

Supplementary materials are available online.

## Implementation and Availability

The methods are implemented in R package `PBtest`, freely and publicly available at https://github.com/yunzhang813/PBtest-R-Package.

