## Supplementary text for "Highly Efficient Hypothesis Testing Methods for Regression-type Tests with Correlated Observations and Heterogeneous Variance Structure"

### S1 Mathematical Derivations

Using the notions introduced in the main manuscript, we first establish some useful identities to facilitate our subsequent mathematical derivations. For $\mathbf{S}$, the *standardized* structure matrix, and $\mathbf{J}$, a matrix of all 1's with corresponding dimensionality, we have

$$\begin{aligned}\mathbf{J}\mathbf{S}^{-1}\mathbf{J} &= \mathbf{1}\left(\mathbf{1}'\mathbf{S}^{-1}\mathbf{1}\right)\mathbf{1}' \\ &= \mathbf{J},\end{aligned} \tag{S1}$$

and

$$\begin{aligned}\left(\mathbf{S}-\mathbf{J}\right)\mathbf{S}^{-1}\left(\mathbf{S}-\mathbf{J}\right) &= (\mathbf{S}-\mathbf{J})(\mathbf{I}-\mathbf{S}^{-1}\mathbf{J}) \\ &= \mathbf{S}-\mathbf{J}-\mathbf{J}+\mathbf{J}\mathbf{S}^{-1}\mathbf{J} \\ &= \mathbf{S}-\mathbf{J}.\end{aligned} \tag{S2}$$

#### S1.1 Centered data

First, we use the generalized least squares approach to estimate the weighted mean for the original data.

$$\begin{aligned}\hat{\mu}(\mathbf{Y}) &= \left(\mathbf{1}'\Sigma^{-1}\mathbf{1}\right)^{-1}\mathbf{1}'\Sigma^{-1}\mathbf{Y} \\ &= \left(\mathbf{1}'\mathbf{S}^{-1}\mathbf{1}\right)^{-1}\mathbf{1}'\mathbf{S}^{-1}\mathbf{Y} \\ &= \mathbf{1}'\mathbf{S}^{-1}\mathbf{Y}.\end{aligned}$$

The last equality is based on Equation (5) in the main manuscript.

Recall that we define the centered data as

$$\mathbf{Y}^{(1)} := \mathbf{Y} - \mathbf{1}\hat{\mu} = \left(\mathbf{I}-\mathbf{J}\mathbf{S}^{-1}\right)\mathbf{Y}.$$

By construction, $\hat{\mu}$ is the best linear unbiased estimator (BLUE) of $\mu$ under $H_0$. Therefore, the joint distribution of $\mathbf{Y}^{(1)}$ under $H_0$ is $\mathbf{Y}^{(1)} \sim \mathcal{N}\left(\mathbf{0},\ \Sigma^{(1)}\right)$, where

$$\begin{aligned}\Sigma^{(1)} &= \left(\mathbf{I}-\mathbf{J}\mathbf{S}^{-1}\right)\sigma^2\mathbf{S}\left(\mathbf{I}-\mathbf{J}\mathbf{S}^{-1}\right)' \\ &= \sigma^2\left(\mathbf{S}-2\mathbf{J}+\mathbf{J}\mathbf{S}^{-1}\mathbf{J}\right) \\ &= \sigma^2\left(\mathbf{S}-\mathbf{J}\right).\end{aligned}$$

*Department of Biostatistics and Computational Biology, University of Rochester, Rochester, New York, U.S.A. Tel: 001-585-275-0666;.

Under $H_1$, $\mathbf{Y}^{(1)}$ has the same covariance matrix and the following expectation vector

$$\begin{aligned}\mathbb{E}(\mathbf{Y}^{(1)}) &= \left(\mathbf{I} - \mathbf{J}\mathbf{S}^{-1}\right)\mathbb{E}(\mathbf{Y})\\ &= \left(\mathbf{I} - \mathbf{J}\mathbf{S}^{-1}\right)(\mathbf{1}\mu + \mathbf{x}\beta)\\ &= \left(\mathbf{I} - \mathbf{J}\mathbf{S}^{-1}\right)\mathbf{x}\beta.\end{aligned}$$

#### S1.2 The B-map

Based on the eigen-decomposition of $\mathbf{S} - \mathbf{J}$ (Equation (6)), we construct $\mathbf{Y}^{(2)}$ as follows

$$\begin{aligned}\mathbf{Y}^{(2)} &:= \Lambda_{n-1}^{-1/2}\mathbf{T}'_{n-1}\mathbf{Y}^{(1)} = \Lambda_{n-1}^{-1/2}\mathbf{T}'_{n-1}\left(\mathbf{I} - \mathbf{J}\mathbf{S}^{-1}\right)\mathbf{Y}\\ &= \Lambda_{n-1}^{-1/2}\mathbf{T}'_{n-1}(\mathbf{S} - \mathbf{J})\,\mathbf{S}^{-1}\mathbf{Y}\\ &= \underbrace{\Lambda_{n-1}^{1/2}\mathbf{T}'_{n-1}\mathbf{S}^{-1}}_{\mathbf{B}}\mathbf{Y} := \mathbf{B}\mathbf{Y} \in \mathbb{R}^{n-1}.\end{aligned}$$

Note that $\mathbf{B}$ is a linear map that maps from the $\mathbb{R}^n$ space to an $\mathbb{R}^{n-1}$ subspace.

#### S1.3 Proof of Theorem 2.1

We first start with a geometric discussion of the operator $\mathbf{P}$. Let $L_{\mathbf{z}}$ be the linear subspace that is spanned by vectors $\mathbf{1}$ and $\mathbf{z}$. By construction, columns in matrix $\mathbf{Q}$ are the orthogonal basis of $L_{\mathbf{z}}$. Matrix $\mathbf{P}$ consists of two operations: $\mathbf{I} - \mathbf{Q}\mathbf{Q}'$ and $\mathbf{Q}\,\mathtt{Rot}\,\mathbf{Q}'$. Note that $\mathbf{Q}\mathbf{Q}'$ is an orthogonal projection onto the subspace $L_{\mathbf{z}}$, so the first operation $\mathbf{I} - \mathbf{Q}\mathbf{Q}'$ is the "complementary" projection on $L_{\mathbf{z}}^{\perp}$, the subspace that is orthogonal to $L_{\mathbf{z}}$. Such a projection will keep any vector in $L_{\mathbf{z}}^{\perp}$ invariant, which is Equation (11). The second operation is a rotation in $L_{\mathbf{z}}$, that rotates vector $\mathbf{z}$ to vector $\mathbf{1}$ by construction, which is Equation (10). Consider an arbitrary vector $\mathbf{v} \in \mathbb{R}^{n-1}$, we can always decompose it into $\mathbf{v} = \mathbf{v}_1 + \mathbf{v}_2$, such that $\mathbf{v}_1 \in L_{\mathbf{z}}$ and $\mathbf{v}_2 \perp \mathbf{v}_1$. The $\mathbf{P}$ transformation will rotate $\mathbf{v}_1$ and keep $\mathbf{v}_2$ invariant; therefore $\mathbf{P}$ must be a orthogonal transformation, which is Equation (9).

Below we present a rigorous proof of Theorem 2.1 based on matrix algebra.

*Proof.* First, let us show that $\mathbf{P}$ is an orthogonal matrix. Remember that $\mathtt{Rot}$ is a rotation matrix such that $\mathtt{Rot}\cdot\mathtt{Rot}' = \mathbf{I}_{2\times 2}$, and $\mathbf{Q}_{(n-1)\times 2}$ is a semi-orthogonal matrix such that $\mathbf{Q}'\mathbf{Q} = \mathbf{I}_{2\times 2}$. We have

$$\begin{aligned}\mathbf{P}\mathbf{P}' &= \left(I - \mathbf{Q}\mathbf{Q}' + \mathbf{Q}\cdot\mathtt{Rot}\cdot\mathbf{Q}'\right)\left(I - \mathbf{Q}\mathbf{Q}' + \mathbf{Q}\cdot\mathtt{Rot}'\cdot\mathbf{Q}'\right)\\ &= \mathbf{I} - \mathbf{Q}\mathbf{Q}' + \mathbf{Q}\cdot\mathtt{Rot}'\cdot\mathbf{Q}' - \mathbf{Q}\mathbf{Q}' + \mathbf{Q}\mathbf{Q}'\mathbf{Q}\mathbf{Q}' - \mathbf{Q}\mathbf{Q}'\mathbf{Q}\cdot\mathtt{Rot}'\cdot\mathbf{Q}'\\ &\quad + \mathbf{Q}\cdot\mathtt{Rot}\cdot\mathbf{Q}' - \mathbf{Q}\cdot\mathtt{Rot}\cdot\mathbf{Q}'\mathbf{Q}\mathbf{Q}' + \mathbf{Q}\cdot\mathtt{Rot}\cdot\mathbf{Q}'\mathbf{Q}\cdot\mathtt{Rot}'\cdot\mathbf{Q}'\\ &= \mathbf{I}.\end{aligned}$$

Next, let us prove Equation (11). By construction, $\mathbf{u}$ is orthogonal to both $\mathbf{1}$ and $\mathbf{z}$. Therefore, $\mathbf{A}'\mathbf{u} = \mathbf{0}$, which implies that $\mathbf{Q}'\mathbf{u} = \mathbf{0}$, based on the assumption that $\mathbf{A}$ is of full-rank. If $\mathbf{A}_{(n-1)\times 2} := (\mathbf{1}_{n-1}|\mathbf{z})$ is not of full-rank, $\mathbf{c}$ must be already in the direction of $\mathbf{1}$ and we don't need the $\mathbf{P}$ transformation in the first place. Hence, $\mathbf{P}\mathbf{u} = (\mathbf{I} - \mathbf{Q}\mathbf{Q}' + \mathbf{Q}\cdot\mathtt{Rot}\cdot\mathbf{Q}')\,\mathbf{u} = \mathbf{u}$.

Lastly, the proof of (10) is as follows. Notice that by definition of QR decomposition, we can write

$$(\mathbf{1}|\mathbf{z}) = (Q_1|Q_2)\begin{pmatrix} R_{11} & R_{12}\\ 0 & R_{22}\end{pmatrix},$$

which implies

$$Q_1 = \frac{\mathbf{1}}{\sqrt{n-1}}, \quad R_{11} = \sqrt{n-1}.$$

Furthermore,

$$\mathbf{z} = Q \begin{pmatrix} R_{12} \\ R_{22} \end{pmatrix} = R_{12}Q_1 + R_{22}Q_2$$

implies that

$$R_{12} = \frac{\langle \mathbf{z},\ \mathbf{1} \rangle}{\sqrt{n-1}} = \xi \cdot \|\mathbf{z}\|, \quad R_{22} = \sqrt{\|\mathbf{z}\|^2 - R_{12}^2} = \sqrt{1-\xi^2} \cdot \|\mathbf{z}\|.$$

Based on the following matrix calculations, we conclude that

$$\begin{aligned}
(\mathbf{I} - \mathbf{Q}\mathbf{Q}')\mathbf{c} &= (\mathbf{I} - \mathbf{Q}\mathbf{Q}')\mathbf{Q} \begin{pmatrix} R_{12} \\ R_{22} \end{pmatrix} = \mathbf{0}, \\
\mathbf{Q} \cdot \texttt{Rot} \cdot \mathbf{Q}'\mathbf{z} &= \mathbf{Q} \cdot \begin{pmatrix} \xi & \sqrt{1-\xi^2} \\ -\sqrt{1-\xi^2} & \xi \end{pmatrix} \cdot \begin{pmatrix} R_{12} \\ R_{22} \end{pmatrix} \\
&= (Q_1 | Q_2) \begin{pmatrix} \boxed{\xi \quad \sqrt{1-\xi^2}} \\ -\sqrt{1-\xi^2} \quad \xi \end{pmatrix} \boxed{\begin{pmatrix} \xi \\ \sqrt{1-\xi^2} \end{pmatrix}} \cdot \|\mathbf{z}\| \\
&= Q_1 \cdot \|\mathbf{z}\| = \frac{\|\mathbf{z}\|}{\sqrt{n-1}} \cdot \mathbf{1}.
\end{aligned}$$

□

Careful readers may notice that there are several choices needed to be made when constructing $\mathbf{P}$. For example, whether to use a clockwise or anti-clockwise rotation matrix, or whether to take positive or negative signs in some of the steps in the QR decomposition. These choices are irrelevant for two-sided tests; but they can affect the direction of one-sided tests.

In our implementation, we force the first row of Rot and the second column of $\mathbf{R}$ (highlighted in red boxes) be identical. In this way, $\mathbf{Q} \cdot \texttt{Rot} \cdot \mathbf{Q}'\mathbf{z} = Q_1 \cdot \|\mathbf{z}\|$ is guaranteed, and the rotation direction and the signs can be determined solely by $Q_1$. As mentioned before, we need to ensure that $\mathbf{P}$ rotates the coefficient vector $\mathbf{z}$ to the direction of $+\mathbf{1}$, so that the one-sided test after rotation is consistent with the original problem. Hence, if $Q_1$ is negative, we will replace $\mathbf{P}$ by $-\mathbf{P}$.

#### S1.4 Proof of Proposition 2.3

*Proof.* We claim that the linear transformation $\mathbf{D}$ and the two constants $u, v$ in Equation (15) can be constructed as follows

$$\begin{aligned}
\mathbf{D} &= \mathbf{T}_{n\times(n-1)} \Lambda^{1/2}_{(n-1)\times(n-1)} \mathbf{P}^{-1}_{(n-1)\times(n-1)}, \\
u &= \frac{\beta\|\mathbf{z}\|}{\sqrt{n-1}}, \qquad v = \mu + \mathbf{1}_n' \mathbf{S}^{-1} \mathbf{x}\beta.
\end{aligned}$$

Here $\mathbf{T}$ and $\Lambda$ are defined in Equation (6), $\mathbf{z}$ is defined in Equation (8), and $\mathbf{P}$ is defined in Theorem 2.1.

Let $\mathbf{Y} := \mathbf{D}\left(\check{\mathbf{Y}} + u\mathbf{1}_{n-1}\right) + (Y^* + v)\mathbf{1}_n$, then

$$
\begin{aligned}
\mathbb{E}(\mathbf{Y}) &= \mathbf{T}\Lambda^{1/2}\left(\mathbf{P}^{-1}\cdot u\mathbf{1}_{n-1}\right) + v\mathbf{1}_n = \mathbf{T}\Lambda^{1/2}\cdot\frac{u\sqrt{n-1}}{\|\mathbf{z}\|}\cdot\mathbf{z} + v\mathbf{1}_n \\
&= \frac{u\sqrt{n-1}}{\|\mathbf{z}\|}\cdot(\mathbf{S}-\mathbf{J})\,\mathbf{S}^{-1}\mathbf{x} + v\mathbf{1}_n \\
&= \beta\left(\mathbf{I} - \mathbf{1}_n\cdot\mathbf{1}_n'\mathbf{S}^{-1}\right)\mathbf{x} + v\mathbf{1}_n \\
&= \beta\mathbf{x} + \left(v - \beta\cdot\mathbf{1}_n'\mathbf{S}^{-1}\mathbf{x}\right)\mathbf{1}_n \\
&= \beta\mathbf{x} + \mu\mathbf{1}_n.
\end{aligned}
$$

And,

$$
\begin{aligned}
\mathrm{cov}(\mathbf{Y}) &= \sigma^2\mathbf{D}\mathbf{D}' + \sigma^2\mathbf{1}_n\mathbf{1}_n' \\
&= \sigma^2\mathbf{T}\Lambda^{1/2}\mathbf{P}^{-1}\cdot\mathbf{P}\Lambda^{1/2}\mathbf{T}' + \sigma^2\mathbf{J} = \sigma^2\mathbf{S}.
\end{aligned}
$$

The second part of Proposition 2.3 claims that the PB-transformation can convert $\mathbf{Y}$ back to a sequence random variables with zero mean if $\beta = 0$ and a nonzero mean otherwise. This is because

$$
\begin{aligned}
&(\mathbf{S}-\mathbf{J})\,\mathbf{S}^{-1}(\mathbf{S}-\mathbf{J}) = \mathbf{S}-\mathbf{J}-\mathbf{J}+\mathbf{J}\mathbf{S}^{-1}\mathbf{J} = \mathbf{S}-\mathbf{J}, \\
&\mathbf{T}\Lambda\mathbf{T}'\cdot\mathbf{S}^{-1}\cdot\mathbf{T}\Lambda\mathbf{T}' = \mathbf{T}\Lambda\mathbf{T}', \qquad \mathbf{T}'\mathbf{S}^{-1}\mathbf{T} = \Lambda^{-1}. \\
\|\mathbf{B}\mathbf{1}_n\|^2 &= \mathbf{1}_n'\mathbf{B}'\mathbf{B}\mathbf{1}_n = \mathbf{1}_n'\mathbf{S}^{-1}\mathbf{T}\Lambda^{1/2}\cdot\Lambda^{1/2}\mathbf{T}'\mathbf{S}^{-1}\mathbf{1}_n \\
&= \mathbf{1}_n'\mathbf{S}^{-1}(\mathbf{S}-\mathbf{J})\,\mathbf{S}^{-1}\mathbf{1}_n = \mathbf{1}_n'\mathbf{S}^{-1}\mathbf{1}_n - \mathbf{1}_n'\mathbf{S}^{-1}\mathbf{J}\mathbf{S}^{-1}\mathbf{1}_n \\
&= 1 - \mathbf{1}_n'\mathbf{S}^{-1}\mathbf{1}_n\mathbf{1}_n'\mathbf{S}^{-1}\mathbf{1}_n = 1 - 1^2 = 0.
\end{aligned}
$$

Therefore,

$$
\begin{aligned}
\mathbf{P}\mathbf{B}\mathbf{D} &= \mathbf{P}\Lambda^{1/2}\mathbf{T}'\mathbf{S}^{-1}\cdot\mathbf{T}\Lambda^{1/2}\mathbf{P}^{-1} = \mathbf{I}_{(n-1)\times(n-1)}, \\
\tilde{\mathbf{Y}} &= \mathbf{P}\mathbf{B}\mathbf{Y} = \mathbf{P}\mathbf{B}\left(\mathbf{D}\left(\check{\mathbf{Y}} + u\mathbf{1}_{n-1}\right) + (Y^*+v)\mathbf{1}_n\right) \\
&= \check{\mathbf{Y}} + u\mathbf{1}_{n-1} + (Y^*+v)\,\mathbf{P}\mathbf{B}\mathbf{1}_n \\
&= \check{\mathbf{Y}} + u\mathbf{1}_{n-1} = \check{\mathbf{Y}} + \frac{\beta\|\mathbf{z}\|}{\sqrt{n-1}}\mathbf{1}_{n-1},
\end{aligned}
$$

Hence, $\mathbb{E}\tilde{\mathbf{Y}} = \mathbf{0}$ if $\beta = 0$ and $\mathbb{E}\tilde{\mathbf{Y}} \neq \mathbf{0}$ otherwise. □

### S2 Correlation Estimation

#### S2.1 Correlation estimation for repeated measurements

In this section, we reframe our regression model in the following matrix form:

$$
\mathbf{y} = \mathbf{X}\boldsymbol{\beta} + \boldsymbol{\epsilon}, \quad \mathbf{X} = (\mathbf{1}, \mathbf{x}), \quad \boldsymbol{\beta} = (\mu, \beta)'.
$$

Suppose there are *a priori* weights available in a diagonal matrix $\mathbf{W}$, define the residuals

$$
\hat{\boldsymbol{\epsilon}}_{\mathrm{WLS}} = \mathbf{y} - \mathbf{X}\hat{\boldsymbol{\beta}}_{\mathrm{WLS}} = \mathbf{y} - \mathbf{X}(\mathbf{X}'\mathbf{W}^{-1}\mathbf{X})^{-1}\mathbf{X}'\mathbf{W}^{-1}\mathbf{y}.
$$

Careful readers may notice that the errors are also correlated due to the repeated measurements. In linear model theory, the above $\hat{\boldsymbol{\beta}}_{\mathrm{WLS}}$ is the BLUE estimator if errors are uncorrelated, and it is an unbiased estimator of $\boldsymbol{\beta}$ if errors are correlated. Therefore, the above residual estimator is valid even with correlated errors in large samples. By construction, the variance-covariance structure of the above residuals preserves the structure in the original data, which consists of the weighting part and the correlation part by design. Before estimating the correlation, we first re-balance the weights and work on the following residual estimates

$$\hat{\boldsymbol{\epsilon}} = \mathbf{W}^{-1}\hat{\boldsymbol{\epsilon}}_{\text{WLS}}. \tag{S3}$$

Further, we have the following assumptions on the correlation structure for repeated measurements:

- Asymptotic normality of the residuals $\hat{\boldsymbol{\epsilon}} = (\hat{\epsilon}_1, \cdots, \hat{\epsilon}_n)'$ such that

  $$\hat{\boldsymbol{\epsilon}} \sim \mathcal{N}(\mathbf{0}, \Sigma_\epsilon).$$

  for some $\Sigma_\epsilon \neq \mathbf{I}$.

- Common variance scaling factor $\sigma^2$, so that

  $$\Sigma_\epsilon = \sigma^2 \cdot \texttt{Cor},$$

  where $\texttt{Cor}$ is the correlation matrix.

- With appropriate data rearrangement, we can write $\texttt{Cor}$ as a block-diagonal matrix. Suppose there are $l = 1, \cdots, L$ blocks,

  $$\texttt{Cor} = \begin{pmatrix} \texttt{Cor}_1 & & \\ & \ddots & \\ & & \texttt{Cor}_L \end{pmatrix}.$$

- Common correlation magnitude $\rho$ for all the blocks, i.e. $\texttt{Cor} = \texttt{Cor}(\rho)$. For each block, assume interchangeable correlation structure, so that

  $$\texttt{Cor}_l(\rho) = \begin{pmatrix} 1 & \rho & \cdots & \rho \\ \rho & 1 & \ddots & \vdots \\ \vdots & \ddots & \ddots & \rho \\ \rho & \cdots & \rho & 1 \end{pmatrix}_{n_l \times n_l} = (1-\rho) \cdot \mathbf{I}_{n_l \times n_l} + \rho \cdot \mathbf{J}_{n_l \times n_l},$$

  where $n_l$ is the size of the $l$th block, and $\mathbf{J}_{n_l \times n_l} = \mathbf{1}_{n_l} \cdot \mathbf{1}'_{n_l}$. Note, $n = \sum_{l=1}^{L} n_l$.

For the above specific structure of the correlation matrix, we derive the following lemma that facilitates our future mathematical computation.

**Lemma S2.1.** *Denote $\mathbf{J} = \mathbf{1}\mathbf{1}'$, the outer product of $\mathbf{1}$ and $\mathbf{1}$. Let $\texttt{Cor} = a\mathbf{I} + b\mathbf{J}$ for $a, b \in \mathbb{R}$ and $\mathbf{I}, \mathbf{J} \in \mathbb{R}^{n \times n}$. Then,*

$$\det(\texttt{Cor}) = a^n\left(1 + \frac{nb}{a}\right) \qquad \textit{and} \qquad \texttt{Cor}^{-1} = \frac{1}{a} \cdot \mathbf{I} - \frac{b}{a(a+nb)} \cdot \mathbf{J}.$$

*Proof.* Write $\texttt{Cor} = \mathbf{E} + \mathbf{u}\mathbf{v}'$ with $\mathbf{E} = a\mathbf{I}, \mathbf{u} = b\mathbf{1}$ and $\mathbf{v} = \mathbf{1}$. The above results follow immediately from the Sherman-Morrison formula. □

##### S2.1.1 The Log-likelihood Function

From the above assumptions, we can work on each block structure separately. For the $l$th block, denote $\boldsymbol{\epsilon}_l$ and $\Sigma_l$ for the corresponding elements, and the log-likelihood for this block is

$$\text{loglik}(\boldsymbol{\epsilon}_l) = -\frac{1}{2}\log\left(\det(\Sigma_l)\right) - \frac{1}{2}\boldsymbol{\epsilon}_l'\Sigma_l^{-1}\boldsymbol{\epsilon}_l + \text{constant}.$$

By Lemma S2.1,

$$\det(\Sigma_l) = \det(\sigma^2 \cdot \texttt{Cor}_l) = \sigma^{2n_l}(1-\rho)^{n_l-1}(1-\rho+n_l\rho),$$
$$\Sigma_l^{-1} = \frac{1}{\sigma^2}\cdot\texttt{Cor}_l^{-1} = \frac{1}{\sigma^2}\left(\frac{1}{1-\rho}\cdot\mathbf{I} - \frac{\rho}{(1-\rho)(1-\rho+n_l\rho)}\cdot\mathbf{J}\right).$$

So,

$$\begin{aligned}-2\cdot\text{loglik}(\boldsymbol{\epsilon}_l) &= n_l\log(\sigma^2) + \log\left((1-\rho)^{n_l-1}(1-\rho+n_l\rho)\right) \\ &\quad + \frac{1}{\sigma^2}\cdot\boldsymbol{\epsilon}_l'\left(\frac{1}{1-\rho}\cdot\mathbf{I} - \frac{\rho}{(1-\rho)(1-\rho+n_l\rho)}\cdot\mathbf{J}\right)\boldsymbol{\epsilon}_l.\end{aligned}$$

By construction, the full log-likelihood function is

$$\mathcal{L}(\rho,\sigma^2) = \text{loglik}(\boldsymbol{\epsilon}) = \sum_{l=1}^{L}\text{loglik}(\boldsymbol{\epsilon}_l).$$

And the maximum likelihood estimators (MLEs) are

$$(\hat{\rho}_{\text{MLE}}, \hat{\sigma}^2_{\text{MLE}}) = \underset{\rho,\sigma^2}{\text{argmax}} -2\mathcal{L}(\rho,\sigma^2).$$

In the following subsections, we are going to derive explicitly the MLEs under the balanced design, i.e. $n_1 = \cdots = n_L = n/L$. Under the unbalanced design, i.e. $n_l$'s are not all the same for $l = 1,\cdots,L$, it is not straightforward to derive explicitly the MLEs. Though gradient descent algorithms may provide numerical solutions to the MLEs, we are going to provide a computationally efficient alternative method based on moment matching.

##### S2.1.2 Balanced Design

WLOG, denote $n_l = n_1 = n/L$ for all $l$. Then

$$\begin{aligned}-2\mathcal{L}(\rho,\sigma^2) &= n\log(\sigma^2) + L\log\left((1-\rho)^{n_1-1}(1-\rho+n_1\rho)\right) \\ &\quad + \frac{1}{\sigma^2}\cdot\underbrace{\sum_{l=1}^{L}\hat{\boldsymbol{\epsilon}}_l'\left(\frac{1}{1-\rho}\cdot\mathbf{I} - \frac{\rho}{(1-\rho)(1-\rho+n_1\rho)}\cdot\mathbf{J}\right)\hat{\boldsymbol{\epsilon}}_l}_{=\left(\frac{1}{1-\rho}SS_1 - \frac{\rho}{(1-\rho)(1-\rho+n_1\rho)}SS_2\right)},\end{aligned}$$

where $SS_1 = \sum_l \hat{\boldsymbol{\epsilon}}_l'\mathbf{I}\hat{\boldsymbol{\epsilon}}_l = \sum_l\sum_i \hat{\epsilon}_{li}^2$ and $SS_2 = \sum_l \hat{\boldsymbol{\epsilon}}_l'\mathbf{J}\hat{\boldsymbol{\epsilon}}_l = \sum_l\sum_{i,j}\hat{\epsilon}_{li}\hat{\epsilon}_{lj}$ are two sufficient statistics.

Take partial derivative with respect to $\sigma^2$ and set to zero,

$$\frac{\partial}{\partial\sigma^2}\left(-2\mathcal{L}(\rho,\sigma^2)\right) = \frac{n}{\sigma^2} - \frac{1}{\sigma^4}\left(\frac{1}{1-\rho}SS_1 - \frac{\rho}{(1-\rho)(1-\rho+n_1\rho)}SS_2\right) = 0$$
$$\Rightarrow \hat{\sigma}^2_{\text{MLE}} = \frac{1}{n}\left(\frac{1}{1-\rho}SS_1 - \frac{\rho}{(1-\rho)(1-\rho+n_1\rho)}SS_2\right).$$

Also, take partial derivative with respect to $\rho$ and set to zero,

$$\frac{\partial}{\partial \rho}\left(-2\mathcal{L}(\rho,\sigma^2)\right) = L\frac{-n_1(n_1-1)\rho}{(1-\rho)(1-\rho+n_1\rho)} + \frac{1}{\sigma^2}\left(\frac{1}{(1-\rho)^2}SS_1 - \frac{1-\rho^2-n_1\rho^2}{(1-\rho)^2(1-\rho+n_1\rho)^2}SS_2\right) = 0.$$

By substituting $\sigma^2 = \hat{\sigma}^2_{\text{MLE}}$ from above, we get

$$\hat{\rho}_{\text{MLE}} = \frac{SS_2 - SS_1}{(n_1-1)SS_1}.$$

###### S2.1.3 Unbalanced Design

If the blocks are not of the same size, i.e. $n_l$'s may be different, then the full log-likelihood becomes

$$-2\mathcal{L}(\rho,\sigma^2) = n\log(\sigma^2) + \sum_{l=1}^{L}\log\left((1-\rho)^{n_l-1}(1-\rho+n_l\rho)\right) + \frac{1}{\sigma^2}\cdot\sum_{l=1}^{L}\hat{\boldsymbol{\epsilon}}_l'\left(\frac{1}{1-\rho}\cdot\mathbf{I} - \frac{\rho}{(1-\rho)(1-\rho+n_l\rho)}\cdot\mathbf{J}\right)\hat{\boldsymbol{\epsilon}}_l.$$

Due to the different $n_l$'s in the denominator, $\hat{\rho}_{\text{MLE}}$ has no explicit form in terms of $\hat{\boldsymbol{\epsilon}}$. Although, we may apply some gradient descent algorithm to obtain the numerical value at convergence, the method is time consuming and may fail to convergent. Here, we provide an alternative method that is computationally efficient and has no convergence issue. By matching the following moments, we have

$$\mathbb{E}SS_1 = n\sigma^2,$$
$$\mathbb{E}(SS_2 - SS_1) = \sigma^2\rho\sum_{l=1}^{L}n_l(n_l-1),$$

where $SS_1$ and $SS_2$ are defined above. So,

$$\hat{\sigma}^2_{\text{moment}} = \frac{SS_1}{n},$$
$$\hat{\rho}^2_{\text{moment}} = \frac{SS_2 - SS_1}{\frac{\sum_{l=1}^{L}n_l(n_l-1)}{n}SS_1}.$$

It is easy to see that $\hat{\rho}^2_{\text{moment}} = \hat{\rho}^2_{\text{MLE}}$ if balanced design.

##### S2.2 Correlation estimation in simulations

The following table compares the correlation estimation based on Equation (13), which is $\hat{\rho}_{\text{PB}}$, or Equation (14), which is $\hat{\rho}_{\text{LMER}}$, in simulation studies. Overall, we find that our proposed correlation estimator has both smaller bias and smaller variance than its counterpart in LMER.

Table S1: Correlation estimation in simulation studies, mean (standard deviation)

| | *SIM1: Group Comparison* | | *SIM2: Regression Test* | |
|---|---|---|---|---|
| | $\rho$=0.2 | $\rho$=0.8 | $\rho$=0.2 | $\rho$=0.8 |
| $\hat{\rho}_{\text{PB}}$ | 0.191 (0.216) | 0.774 (0.096) | 0.179 (0.176) | 0.768 (0.124) |
| $\hat{\rho}_{\text{LMER}}$ | 0.326 (0.340) | 0.887 (0.178) | 0.274 (0.213) | 0.847 (0.075) |

### S3 Simulation Design

In our simulation studies, we design the covariance structure of an $n$-dimensional joint distribution based on Equation (12) in the main manuscript. Let $\mathbf{W} = \text{diag}(w_1, \cdots, w_n)$, and the weights are $w_i = 1/\sigma_i^2$ for $i = 1, \cdots, n$, where $\sigma_i^2$ is the variance of the $i$th marginal distribution. We specify `Cor` in the following two simulation scenarios.

SIM1: This simulation is designed for completely paired group comparison. Suppose there are two groups, namely A and B. The total sample size is $n = 40$, within which $n_{\text{A}} = n_{\text{B}} = 20$. All observations are paired in the two groups. The correlation matrix for SIM1 can be expressed as

$$\texttt{Cor}(\rho) = \begin{pmatrix} \begin{smallmatrix} 1 & \rho \\ \rho & 1 \end{smallmatrix} & & \\ & \ddots & \\ & & \begin{smallmatrix} 1 & \rho \\ \rho & 1 \end{smallmatrix} \end{pmatrix}.$$

The corresponding group indicator is $\mathbf{1}_{\text{B}} = (0, 1, \cdots, 0, 1)'$. The weights, $\sigma_1^2, \cdots, \sigma_n^2$, are randomly drawn from Unif$(1, 6)$.

SIM2: This simulation is designed for regression-type test with a continuous covariate. Let $n = 80$ and generate the covariate from $N(3, 1)$. The observations are mixed with singletons, doubles, and triplets, i.e. subjects with one, two, and three repeated measurements, respectively. Specifically, we generate 10 doubles and 10 triplets; and the rest are singletons. The correlation matrix for SIM2 can be expressed as

$$\texttt{Cor}(\rho) = \begin{pmatrix} 1 & & & & & \\ & \ddots & & & & \\ & & \begin{smallmatrix} 1 & \rho \\ \rho & 1 \end{smallmatrix} & & & \\ & & & \ddots & & \\ & & & & \begin{smallmatrix} 1 & \rho & \rho \\ \rho & 1 & \rho \\ \rho & \rho & 1 \end{smallmatrix} & \\ & & & & & \ddots \end{pmatrix}.$$

For the weights, $\sigma_1^2, \cdots, \sigma_n^2$ are random draws from Unif$(1, 2)$.

For each simulation scenario, we consider two underlying distributions, i.e. normal distribution and double exponential distribution; as well as two correlation levels, i.e. $\rho = 0.2$ and $\rho = 0.8$. We generate the correlated observations using Proposition 2.3. In each simulation, we test 2,000 hypotheses simultaneously: 1,000 from the null distribution and 1,000 from the alternative distribution. We repeat 10 times for each simulation.

In the simulation studies, we investigated two versions of the PB-transformed tests, i.e. with $\rho$ (oracle) or $\hat{\rho}$ (estimation), and compared the proposed parametric and semiparametric tests to the following methods in each category.

- SIM1 & normal: weighted linear mixed-effects model with test for the fixed effect (weighted LMER), paired $t$-test, weighted regression $t$-test, and two-sample $t$-test.
- SIM1 & double exponential: test performed by `svyranktest()`, Friedman test, Wilcoxson signed rank test, and Wilcoxon rank-sum test.
- SIM2 & normal: weighted LMER, weighted regression $t$-test, and two-sample $t$-test.
- SIM2 & double exponential: Spearman rank correlation tests with and without B-transformation. One important purpose of this simulation is to show the advantage of the P-map.

To our best knowledge, the above tests are the most widely used practical solutions for each category.

### S4 Real Data Analyses

In this study, we illustrate the PB-transformed tests in the context of differential expression analysis for next-generation sequencing data.

RNA-seq (Wang et al., 2009) is a deep-sequencing technology that is widely used in gene expression experiments. Depending on different experiment choices, e.g. instrument platforms, sequencing libraries preparation, etc., the number of reads sequenced per sample varies largely ($10^7 - 10^{12}$ (Sims et al., 2014)) from study to study. In addition to the technical variation, the number of read sequenced is also strongly related with the true biological variation within the specimen that has been sampled. Formally, *sequencing depth* is defined as the total number of reads sequenced in each sample. Considerations of sequencing depth in genomic analyses are reviewed in Sims et al. (2014). Here, we use the PB-transformed test to adjust the varying sequencing depth in RNA-seq data analyses.

We download RNA-seq data of breast invasive carcinoma from The Cancer Genome Atlas (TCGA) (Cancer Genome Atlas Network, 2012). Primarily, breast cancers can be classified into subtypes based on human epidermal growth factor receptor 2 (HER2) gene (Onitilo et al., 2009; Carey et al., 2006). In this study, we select 35 samples from tissue source site "BH", which have the following clinical traits: HER2-positive, white race and female sex. In this tissue source site, a subset of the tumor specimens have companion normal tissue specimens.

#### S4.1 Data preprocessing

Raw RNA-seq count data are normalized by reads per kilobase million (RPKM) (Mortazavi et al., 2008). Figure S1 shows data preprocessing criteria. Sample filtering is based on between-sample correlations (Figure S1a). Samples with very low similarities with other samples, which is defined as median of $(\text{corr}_{i,j})_j \leq 0.6$, where $\text{corr}_{i,j}$ is the correlation between sample $i$ and sample $j$, are removed. After this step, 28 samples are kept; and the mean correlation across these remaining samples is 0.72.

Next, we apply nonspecific filtering based on log2-transformed gene expression values (Figure S1b). The log2-transformation is defined as $\log_2(X + 1)$, where $X$ are the normalized counts. This histogram shows that the distribution of mean expression level for individual genes can be roughly divided into "low" and "high" expression clusters. Low expression genes (i.e. mean expression value after log2-transformation is $\leq 1.5$) are filtered, and 11,453 genes remain in the working data.

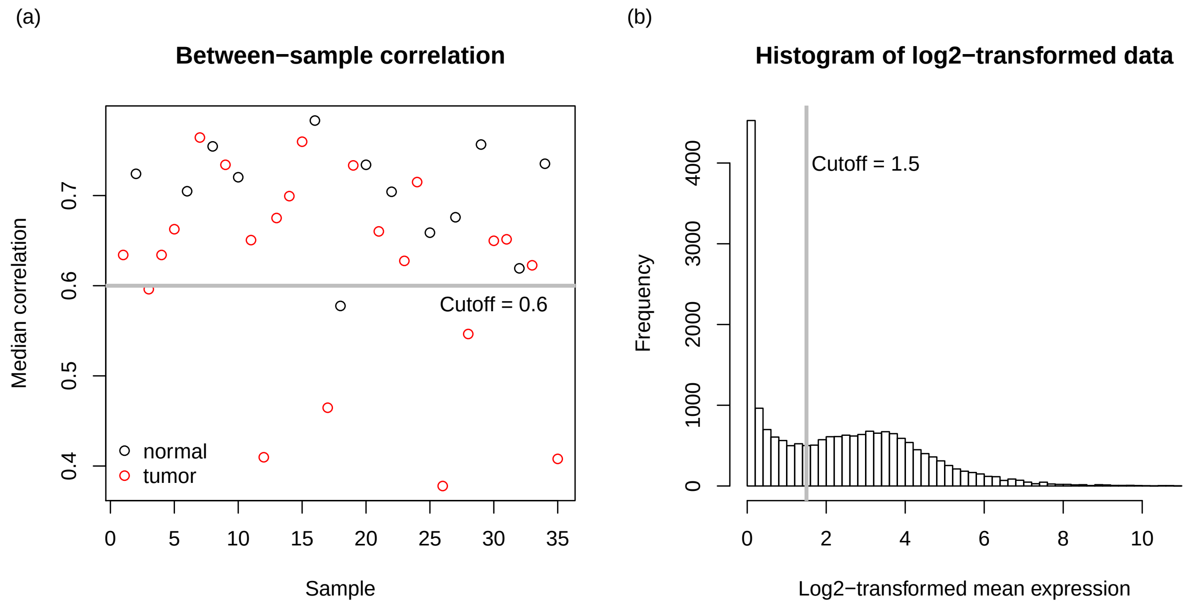


Figure S1: Criteria of RNA-seq data preprocessing.

#### S4.2 Weights for RNA-seq samples

To justify the choice of weights in Equation (18), we consider a simple count model based on the binomial distribution. The same principle is extendable to more advanced models. Borrowing the notations in Robinson et al. (2010), let $Y_{ij}$ be the raw count for gene $i$ in sample $j$. We assume that $Y_{ij} \sim \text{Binom}(M_j, p_i)$, where $M_j$ is the sequencing depth (or library size), and $p_i$ is the relative abundance of gene $i$. The null hypothesis is $H_0 : p_{i,\text{tumor}} = p_{i,\text{normal}}$. By large sample theory,

$$\sqrt{M_j}\,(\hat{p}_i - p_i) \to \text{N}(0, \sigma^2), \tag{S4}$$

where $\sigma^2$ is the variance for the binomial distribution. Equation (S4) shows that the sampling variability of the relative abundance $p_i$ is inversely proportional to $M_j$. The same large sample property can be applied to more complex models such as Negative Binomial model (Robinson & Smyth, 2007; Anders & Huber, 2010; Robinson et al., 2010; Love et al., 2014).

#### S4.3 Criteria for differentially expressed gene (DEG)

In our real data analyses, Benjamini-Hochberg procedure (Benjamini & Hochberg, 1995) is applied to adjust the raw $p$-values so that we can control the false discovery rate (FDR) at 0.05 level. We also threshold the fold change (FC), which is defined as

$$\text{FC}_i = \frac{\sum_{j=1}^{n_A} x_{ij}/n_A}{\sum_{j=1}^{n_B} y_{ij}/n_B} \quad \text{for gene } i,$$

where $x_{ij}$ and $y_{ij}$ are expression values, and $n_A$ and $n_B$ are sample sizes for the tumor and normal groups, respectively. FC is a popular gene selection criterion among practitioners, as it reflects the actual mean expression difference between the groups. Using both criteria, we define DEG as the gene with BH-adjusted $p$-value $< 0.01$, and FC $< 1/2$ (downregulated) or $> 2$ (upregulated).

##### S4.4 Biology of DEG lists

The eleven genes uniquely identified by the PB-transformed *t*-test are known to be involved in cell survival, proliferation and migration. The CXCR4-CXCL12 chemokine signaling pathway is one of the deregulated signaling pathway uniquely identified by PB-transformed t-test in HER2+ breast cancer cells. This pathway is known to play a crucial role in promoting breast cancer metastasis and has been reported to be associated with poor prognosis (Sun et al., 2014; Müller et al., 2001). Metastatic breast tumor cells have been shown to express high levels of CXCR4 (Xu et al., 2015). After binding to its ligand CXCL12, CXCR4 activates downstream PI3K and MAP kinase signaling pathways that regulate cell proliferation, survival and chemotaxis (Xu et al., 2015; Andreou et al., 2012). CXCL12 is constitutively expressed by different cell types in tissues such as lung, liver and bone marrow (Xu et al., 2015; Cojoc et al., 2013). Secreted CXCL12 from these healthy tissues drives metastatic invasion of CXCR4+ tumor cells into normal tissues (Xu et al., 2015; Sun et al., 2010; Sarvaiya et al., 2013). The PB-transformed *t*-test also identifies increased PI3KR2 (the regulatory p85 subunit of PI3K) and MAP kinase 13 (p38 $\delta$-subunit). Hyper-activation of both PI3K p85 and p38 were shown to be associated with different types of metastases (Cuenda & Sanz-Ezquerro, 2017; Martini et al., 2014). Both of these intracellular molecules can be activated by CXCR4 signaling (Xu et al., 2015; Cojoc et al., 2013; Sun et al., 2002). Our results demonstrate that these HER2+ breast cancer cells not only had increased CXCR4 transcripts, but also had elevated transcriptional levels of its downstream signaling molecules to support CXCR4-triggered metastasis. Another critical gene annotated by PB-transformed *t*-test is GAB2. GAB2 has been reported to be upregulated in primary breast tumors as well as in breast cancer cell lines (Bentires-Alj et al., 2006). Co-expression of GAB2 along with HER2 is pivotal to acquire metastatic property by HER2+ cells (Bentires-Alj et al., 2006). Activated GAB2 can in turn stimulate several intracellular signaling molecules such as SHC and p85 subunit of PI3K (Bentires-Alj et al., 2006; Ding et al., 2015), both uniquely annotated by our PB-transformed *t*-test (SHC2 and PIK3R2).

In contrast, the weighted LMER uniquely identifies six important genes from the same set of data. Three of those genes (COX6B1, COX4I2 and COX5A) are different subunits of cytochrome C oxidase. Another gene, UQCR10, is a cytochrome c reductase. These enzymes are essential for ATP generation during mitochondrial respiration. Since tumor cells need more energy than most of the normal cells, it is expected that those breast cancer cells had over-expression of those genes. Another gene identified by LMER is HSP70 family member HSPA5. The HSP70 family proteins are known to transduce anti-apoptotic signals in many types of cancer cells (Murphy, 2013). Multiple HSP70 inhibitors are being tested for anti-tumor activity (Murphy, 2013).

##### S4.5 Functional annotation terms from DAVID gene set analyses

Table S2 shows the functional annotation terms identified by DAVID gene set analysis based on the unique genes exclusively belonging to one of the top 2,000 DEG lists of the PB-transformed *t*-test and the weighted LMER test.

Table S2: Functional annotations based on the unique genes in each of the top 2,000 gene lists.

| | | GO terms or KEGG pathways |
|---|---|---|
| **PB-transformed *t*-test** | | |
| | 1 | GO:0007059~chromosome segregation |
| | 2 | GO:0007155~cell adhesion |
| | 3 | hsa04670:Leukocyte transendothelial migration |

4 hsa04914:Progesterone-mediated oocyte maturation
5 hsa04923:Regulation of lipolysis in adipocytes
6 GO:0030198~extracellular matrix organization
7 GO:0007165~signal transduction
8 hsa04022:cGMP-PKG signaling pathway
9 GO:0018105~peptidyl-serine phosphorylation
10 hsa04666:Fc gamma R-mediated phagocytosis
11 GO:0007204~positive regulation of cytosolic calcium ion concentration
12 hsa04015:Rap1 signaling pathway
13 hsa04810:Regulation of actin cytoskeleton
14 hsa04664:Fc epsilon RI signaling pathway
15 hsa04370:VEGF signaling pathway
16 hsa04510:Focal adhesion
17 hsa05230:Central carbon metabolism in cancer
18 GO:0035556~intracellular signal transduction
19 hsa04062:Chemokine signaling pathway
20 GO:0070527~platelet aggregation
21 GO:0008015~blood circulation
22 hsa04917:Prolactin signaling pathway
23 GO:0051056~regulation of small GTPase mediated signal transduction
24 GO:0050920~regulation of chemotaxis
25 hsa04611:Platelet activation
26 GO:0048015~phosphatidylinositol-mediated signaling
27 hsa05214:Glioma
28 GO:0010715~regulation of extracellular matrix disassembly
29 GO:0031100~organ regeneration
30 hsa05132:Salmonella infection
31 GO:0006110~regulation of glycolytic process
32 hsa04071:Sphingolipid signaling pathway
33 GO:0009615~response to virus
34 GO:0010038~response to metal ion
35 hsa04024:cAMP signaling pathway
36 GO:0006260~DNA replication
37 hsa05220:Chronic myeloid leukemia
38 hsa05100:Bacterial invasion of epithelial cells
39 hsa04014:Ras signaling pathway
40 GO:0019064~fusion of virus membrane with host plasma membrane
41 hsa04068:FoxO signaling pathway

**Weighted LMER**

1 GO:1902600~hydrogen ion transmembrane transport
2 hsa04932:Non-alcoholic fatty liver disease (NAFLD)
3 hsa05010:Alzheimer's disease
4 GO:0006379~mRNA cleavage
5 GO:1990440~positive regulation of transcription from RNA polymerase II promoter in response to endoplasmic reticulum stress
6 hsa04260:Cardiac muscle contraction
7 hsa00190:Oxidative phosphorylation

| | |
|---|---|
| 8 | hsa05016:Huntington's disease |
| 9 | hsa05012:Parkinson's disease |
| 10 | GO:0006123~mitochondrial electron transport, cytochrome c to oxygen |
| 11 | GO:0036498~IRE1-mediated unfolded protein response |
| 12 | GO:0030574~collagen catabolic process |
| 13 | GO:0043065~positive regulation of apoptotic process |
| 14 | GO:2000110~negative regulation of macrophage apoptotic process |
| 15 | GO:0030968~endoplasmic reticulum unfolded protein response |
| 16 | GO:0051603~proteolysis involved in cellular protein catabolic process |
| 17 | GO:0021762~substantia nigra development |
| 18 | GO:0006987~activation of signaling protein activity involved in unfolded protein response |
| 19 | hsa04141:Protein processing in endoplasmic reticulum |
| 20 | GO:0030308~negative regulation of cell growth |
| 21 | hsa04620:Toll-like receptor signaling pathway |
| 22 | GO:0006983~ER overload response |
| 23 | GO:0001501~skeletal system development |
| 24 | GO:0090162~establishment of epithelial cell polarity |
| 25 | GO:0036499~PERK-mediated unfolded protein response |
| 26 | GO:0033138~positive regulation of peptidyl-serine phosphorylation |
| 27 | GO:0071260~cellular response to mechanical stimulus |
| 28 | GO:0098719~sodium ion import across plasma membrane |
